# GPCR surface creates a favorable pathway for membrane permeation of drug molecules

**DOI:** 10.1101/2024.03.18.585530

**Authors:** Cristina Gil Herrero, Sebastian Thallmair

**Affiliations:** Frankfurt Institute for Advanced Studies, Ruth-Moufang-Straße 1, 60438 Frankfurt am Main (Germany); Faculty of Biochemistry, Chemistry and Pharmacy, Goethe University Frankfurt, Frankfurt am Main (Germany)

## Abstract

G protein-coupled receptors (GPCRs) play a crucial role in modulating physiological responses and serve as the main drug target. Specifically, salmeterol and salbutamol which are used for the treatment of pulmonary diseases, exert their effects by activating the GPCR β2-adrenergic receptor (β2AR). In our study, we employed coarse-grained molecular dynamics simulations with the Martini 3 force field to investigate the dynamics of drug molecules in membranes in presence and absence of β2AR. Our simulations reveal that in more than 50% of the *flip-flop* events the drug molecules use the β2AR surface to permeate the membrane. The pathway along the GPCR surface is significantly more energetically favorable for the drug molecules, which was revealed by umbrella sampling simulations along spontaneous *flip-flop* pathways. Furthermore, we assessed the behavior of drugs with intracellular targets, such as kinase inhibitors, whose therapeutic efficacy could benefit from this observation. In summary, our results show that β2AR surface interactions can significantly enhance membrane permeation of drugs, emphasizing their potential for consideration in future drug development strategies.

---

G protein-coupled receptors (GPCRs) are the largest family of integral membrane proteins, which transform extracellular signals into intracellular responses via coupling to guanosine triphosphate (GTP)-binding proteins (G proteins). GPCRs are sensitive to light, taste or odour molecules, neurotransmitters, and hormones triggering the vast majority of physiological responses to these messengers. Thus, they mediate most of the physiological stimuli into cellular reactions [1].

These properties provide them great potential as therapeutic targets. Therefore, it is not surprising that most approved drugs to date target GPCRs, establishing this group as the largest family of protein drug targets [2]. One common feature of GPCRs is their structure, consisting of seven transmembrane domains with an *α*-helical conformation (H1-H7) connected by extracellular (ECL1-ECL3) and intracellular loops (ICL1-ICL3) [3, 4]. Upon receiving an extracellular signal, GPCRs transmit the message to the cell interior via specific conformational changes. The protein changes towards its active conformation through the opening of the structure in the cytoplasmic region, enabling interaction with heterotrimeric G proteins or through G protein independent pathways involving, for instance, arrestin [5].

Although GPCRs are mainly known for their essential roles in signal transduction across the plasma membrane, recent in vitro studies [6, 7, 8] as well as molecular dynamics simulations [9] have revealed a moonlighting activity of these receptors: lipid scrambling. *Flip-flops* are rarely performed spontaneously due to the high energy barrier for the polar lipid headgroup to traverse the region of the non-polar tails. Nevertheless, it is a crucial mechanism in the cell to preserve the asymmetry of the plasma membrane, cell growth, or adaptation of cellular responses to physiological challenges among other functions. Therefore, specialized transporters denominated flippases and floppases, respectively, exist in cells. Some of them hydrolyze ATP to produce a unidirectional transport of lipids against a concentration gradient. Other lipid transporters that are more challenging to identify are scramblases, which typically facilitate the translocation of different lipid types in both directions, as it was found for some GPCRs including opsin, β2-adrenergic receptor (β2AR), and adenosine A2A receptor [10]. The proposed mechanism is denominated card reader mechanism and relies on the interaction of the lipid headgroup with the GPCR. In this mechanism, the lipid headgroup slides along a hydrophilic pathway on the surface of the GPCR through the membrane, while the lipid tails reside in the bilayer core [11]. Atomistic MD simulations revealed a pathway between the transmembrane helices H6 and H7 for opsin which exhibits an experimental *flip-flop* rate of 1/(100 μs). Similar rates were reported also for β1- and β2-adrenergic receptors, as well as the A2A receptor [7, 9].

β2-adrenegic receptor (β2AR) is a well-characterized GPCR located predominantly in the smooth muscle cells of the pulmonary tissue but also in the cardiovascular system [12]. It exhibits the general structure typical of GPCRs and can alternate between active and inactive conformations [13]. Salmeterol (SALMT) and salbutamol (SALBT) (or albuterol) are partial β2AR-agonists employed in the treatment of several respiratory diseases such as asthma and present a high affinity to β2AR. Both drugs have similar chemical structures but exhibit different pharmacological profiles. SALMT is highly soluble in organic solvents resulting in an elevated membrane partitioning, a characteristic that confers it a long-acting behavior as bronchodilator [14]. On the other hand, SALBT has the same polar head as SALMT, consisting of an aromatic ring with two hydrophilic substituents, but lacks the elongated hydrophobic tail of SALMT. Therefore, SALBT is a short-acting β2AR-agonist due to its lower lipophilicity that results in low membrane partitioning [15].

Given the previously discovered role of GPCRs as scramblases, our research is driven by the intriguing possibility that this membrane translocation could be extended to other molecules like drugs. Thus, considering the ubiquity of this class of proteins in the body, drug delivery could benefit from this new mechanism, a new approach that could help treatment methods throughout the human body. To this end, we investigate the transmembrane *flip-flop* of SALMT and SALBT on the β2AR surface using the Martini 3 coarse-grained (CG) force field. After briefly discussing the CG ligand models, we explore the main aspects of their interaction with β2AR relevant for the *flip-flop*. We identify the helices along which most of the *flip-flops* take place and conduct umbrella sampling (US) simulations along spontaneous *flip-flop* pathways to compare the energetics on the protein surface to the pure membrane. Finally, we show that the effect facilitating membrane translocation is not limited to β2AR drug molecules. We also observe the *flip-flop* enhancement for two approved kinase inhibitors – baricitinib and dasatinib – which have to access the cell interior to exert their therapeutic effects.

For the parametrization of SALMT and SALBT, we followed the guidelines for Martini 3 models [16, 17, 18]. The bead type assignment is depicted in Figure 1B and C, where the CG beads are colored according to their size. More details about the bead type assignment, optimization of bonded terms, and model validation are provided in Section 2 of the Supporting Information (SI).

**Figure 1:**
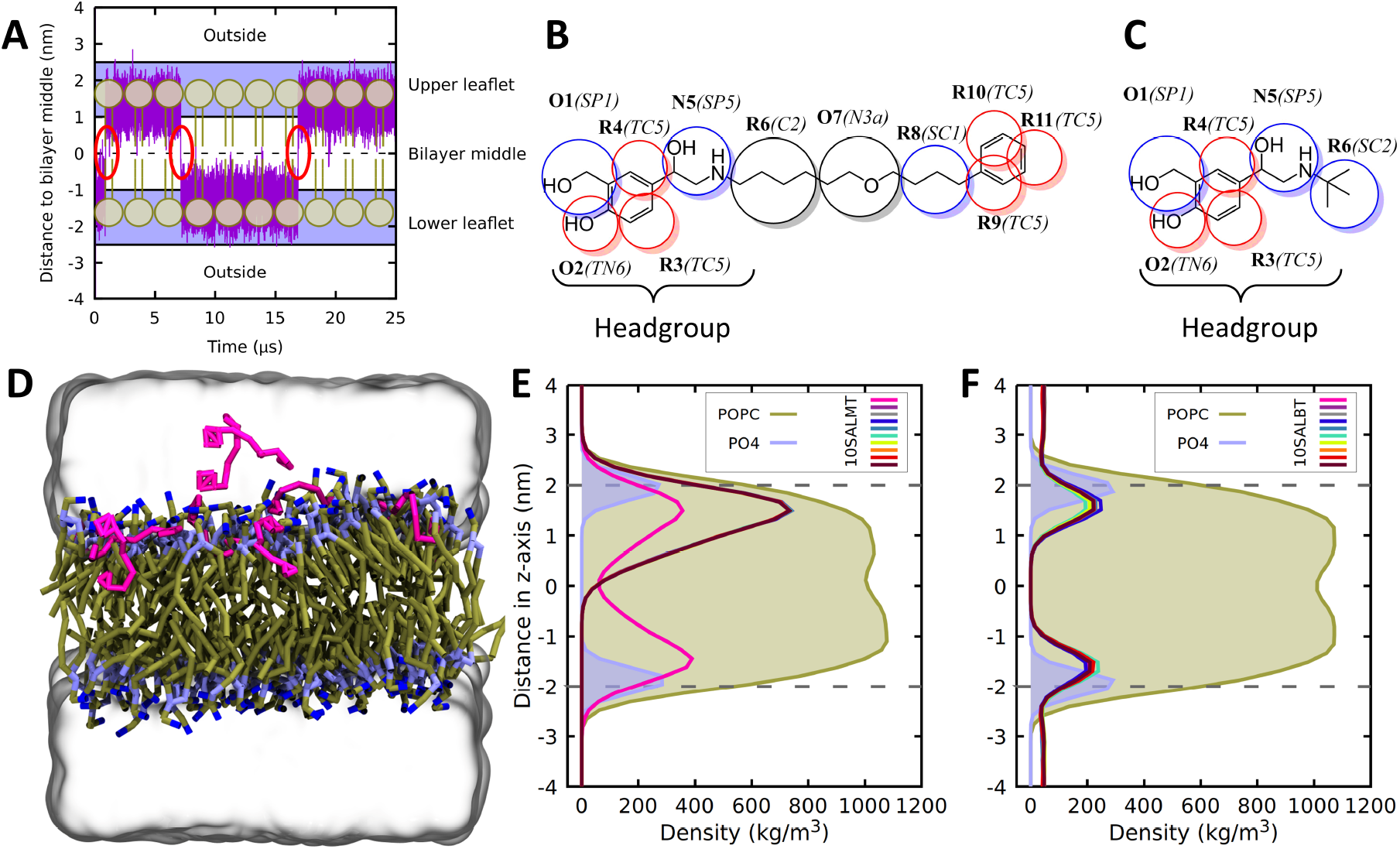
(A) Distance of the center of mass (COM) of a SALMT molecule to the COM of the terminal beads in the lipid tails along a 25 μs simulation trajectory. The location of the lipids is shown schematically as well as the boundaries of the four regions used to count the number of *flip-flops*. (B) CG mapping of SALMT and (C) SALBT. The Martini 3 bead type is given in brackets after the bead name; tiny (T) beads are colored red, small (S) beads blue, and regular (R) beads black. (D) System setup containing ten SALMT molecules and a POPC membrane solvated with water including 0.15 M NaCl. (E) Density distribution of ten SALMT and (F) SALBT molecules, the POPC membrane, and the lipids’ PO4 beads depicted along the membrane normal (z-axis) from a 25 μs simulation trajectory. The intracellular region corresponds to negative z-values; the extracellular region to positive z-values. The densities of SALMT and SALBT are scaled by a factor of 100.

With the optimized drug molecule models at hand, we first investigated their behavior in a lipid membrane. Ten drug molecules were placed above a 1-palmitoyl-2-oleoyl-sn-glycero-3-phosphocholine (POPC) membrane, and five replicas of 25 μs each were run for both SALMT and SALBT, respectively.

In all five replicas, the SALMT molecules quickly entered the membrane. Due to an asymmetric starting distribution in the water phase, they mostly entered one of the two leaflets - the upper one in Figure 1D. The density distributions of each individual SALMT molecule along the membrane normal, the z-axis, are depicted in Figure 1E for one representative replica. The water phase corresponds to the z-range outside the POPC density depicted in green. The SALMT density in the water phase is negligible, which correctly describes its known high lipophilicity. Remarkably, all but one SALMT molecule solely resided in the upper leaflet at positive z values along the membrane normal. This one SALMT (pink solid line) performed a transverse diffusion through the membrane center to the opposite leaflet, a so-called *flip-flop* and spent approximately half of the simulation time in each leaflet. The density distributions for one replica with SALBT are depicted in Figure 1F. In contrast to SALMT, SALBT shows a non-negligible density in the water phase. This indicates a reduced lipophilicity for SALBT in comparison to SALMT. Nevertheless, SALBT molecules also enter the membrane and the maxima of their density distributions match the one of SALMT. Note that, unlike SALMT, the *flip-flops* of SALBT cannot be observed in the density distributions since the drug molecules easily diffuse through the periodic boundary conditions along the z-axis. Hence, they are able to populate the other leaflet without performing any *flip-flop*. Overall, SALMT exhibits a higher lipophilic character than SALBT (Figure 1E and F) in our CG simulations. This is in good agreement with their respective pharmacological characteristics of long-acting (SALMT) and short-acting drugs (SALBT) which is attributed to their difference in lipophilicity [19].

Next, we tested whether the presence of β2AR would impact the drug molecule behavior in the membrane. To this end, we set up a system with ten drug molecules in the solvent phase above a POPC membrane with one β2AR embedded. The simulation setup for SALMT is depicted in Figure 2A. Again, five replicas of 25 μs were simulated with SALMT and SALBT, respectively.

**Figure 2:**
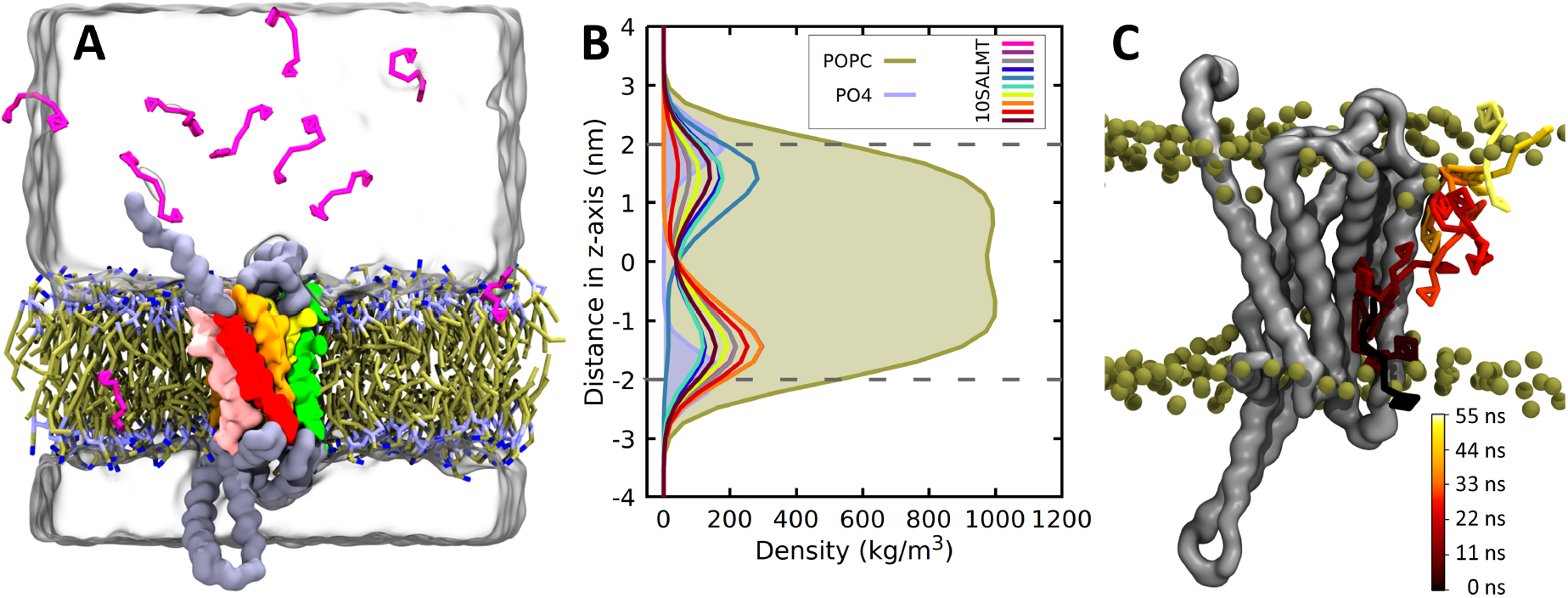
(A) System setup containing ten SALMT molecules and β2AR embedded in a POPC membrane solvated with water including 0.15 M NaCl. (B) Density distribution of ten SALMT molecules, the POPC membrane, and the lipids’ PO4 beads depicted along the membrane normal (z-axis) from a 25 μs simulation trajectory. The intracellular region corresponds to negative z-values; the extracellular region to positive z-values. The densities of the SALMT molecules are scaled by a factor of 100. (C) Snapshots of a *flip-flop* event of SALMT along the protein (grey) surface from the lower to the upper leaflet. After aligning the trajectory to the protein backbone, only one intermediate protein snapshot is depicted while snapshots of the *flip-flopping* SALMT are colored from black to yellow representing the evolution in time. The lipids’ PO4 beads are shown as green spheres indicating the position of the membrane.

In the case of SALMT, the density distributions showed a pronounced difference compared to the simulations in the pure POPC membrane (Figure 2B). Almost all SALMT molecules exhibit densities in both leaflets, which implies that they performed at least one *flip-flop*. Based on these observations, β2AR seems to enhance the *flip-flop* of SALMT. For SALBT, the density distributions are similar to the ones in the absence of β2AR, where leaflet changes are already possible via the water phase because of the periodic boundary conditions (Figure S7). Figure 2C shows a representative *flip-flopping* event of SALMT, where it moves from the lower to the upper leaflet at the surface of β2AR. The time evolution is indicated by the color scale from black to yellow.

In order to quantify the enhancement of *flip-flops* by β2AR, we evaluated the number of *flip-flops* in each of the simulated systems. Table 1 shows the number of *flip-flops* in each direction for both drug molecules with and without β2AR. In the systems with protein, we additionally evaluated the distance between the COMs of the drug molecule and the protein to distinguish between *flip-flops* in the pure membrane and on the protein surface. The total number of *flip-flops* for SALMT and SALBT clearly increases when β2AR is present. The ratio of *flip-flops* on the β2AR surface in the case of SALMT is 18 out of 22 *flip-flops* downwards and 13 out of 24 *flip-flops* upwards. Notably, 67% of the *flip-flop* events take place on the protein surface despite the annular membrane area of β2AR (up to a distance of 1 nm from the protein) being only about 6-7% of the total membrane area. While only 11% of the SALMT molecules are on average in contact with the β2AR surface, they perform more than half of the *flip-flop* events.

**Table 1:** Number of *flip-flops* of SALMT and SALBT calculated for five replicas (25 μs simulation time each) of two different system setups; with and without β2AR.

| SALMT |  |  |  | SALBT |  |  |  |
| --- | --- | --- | --- | --- | --- | --- | --- |
| | Only | β2AR and POPC | | | Only | β2AR and POPC | |
| | POPC | Total | On β2AR | | POPC | Total | On β2AR |
| Rep 1 | 3 ↓ 2 ↑ | 3 ↓ 3 ↑ | 1 ↓ 0 ↑ | Rep 1 | 1 ↓ 2 ↑ | 4 ↓ 4 ↑ | 1 ↓ 2 ↑ |
| Rep 2 | 3 ↓ 0 ↑ | 2 ↓ 5 ↑ | 2 ↓ 3 ↑ | Rep 2 | 1 ↓ 1 ↑ | 3 ↓ 2 ↑ | 3 ↓ 0 ↑ |
| Rep 3 | 1 ↓ 1 ↑ | 6 ↓ 5 ↑ | 5 ↓ 4 ↑ | Rep 3 | 3 ↓ 2 ↑ | 4 ↓ 3 ↑ | 1 ↓ 2 ↑ |
| Rep 4 | 2 ↓ 2 ↑ | 5 ↓ 4 ↑ | 4 ↓ 1 ↑ | Rep 4 | 2 ↓ 1 ↑ | 1 ↓ 0 ↑ | 0 ↓ 0 ↑ |
| Rep 5 | 2 ↓ 1 ↑ | 6 ↓ 7 ↑ | 6 ↓ 5 ↑ | Rep 5 | 3 ↓ 2 ↑ | 2 ↓ 1 ↑ | 2 ↓ 0 ↑ |
| **Total** | 11 ↓ 6 ↑ | 22 ↓ 24 ↑ | 18 ↓ 13 ↑ | **Total** | 10 ↓ 8 ↑ | 14 ↓ 10 ↑ | 7 ↓ 4 ↑ |

The ratio of *flip-flops* on the β2AR surface is lower in the case of SALBT, with 7 out of 14 *flip-flops* downwards and 4 out of 10 *flip-flops* upwards. However, still 46% of them take place on the protein surface. The overall reduced number of *flip-flops* is due to the reduced concentration of SALBT in the membrane, as indicated by our test simulations with a flat-bottom potential (see Table S2 in the SI). In particular, only 7% of the SALBT molecules are in contact with the protein surface on average.

To gain more insights about where on the β2AR surface *flip-flops* occur, we calculated the number of contacts between the individual residues and the polar drug molecule heads using a distance cutoff of 0.7 nm. The results are shown in Figure 3A and B for SALMT and SALBT, respectively. The colored rectangles indicate the different transmembrane helices (H1-H7) of β2AR. For both drug molecules, oscillations in the number of contacts appear due to subsequent residues in an α-helix located alternately on the protein surface and in its interior. Beyond these oscillations, more contacts are observed at the water-membrane interface, i.e. the respective borders of the shaded regions, compared to the bilayer middle.

**Figure 3:**
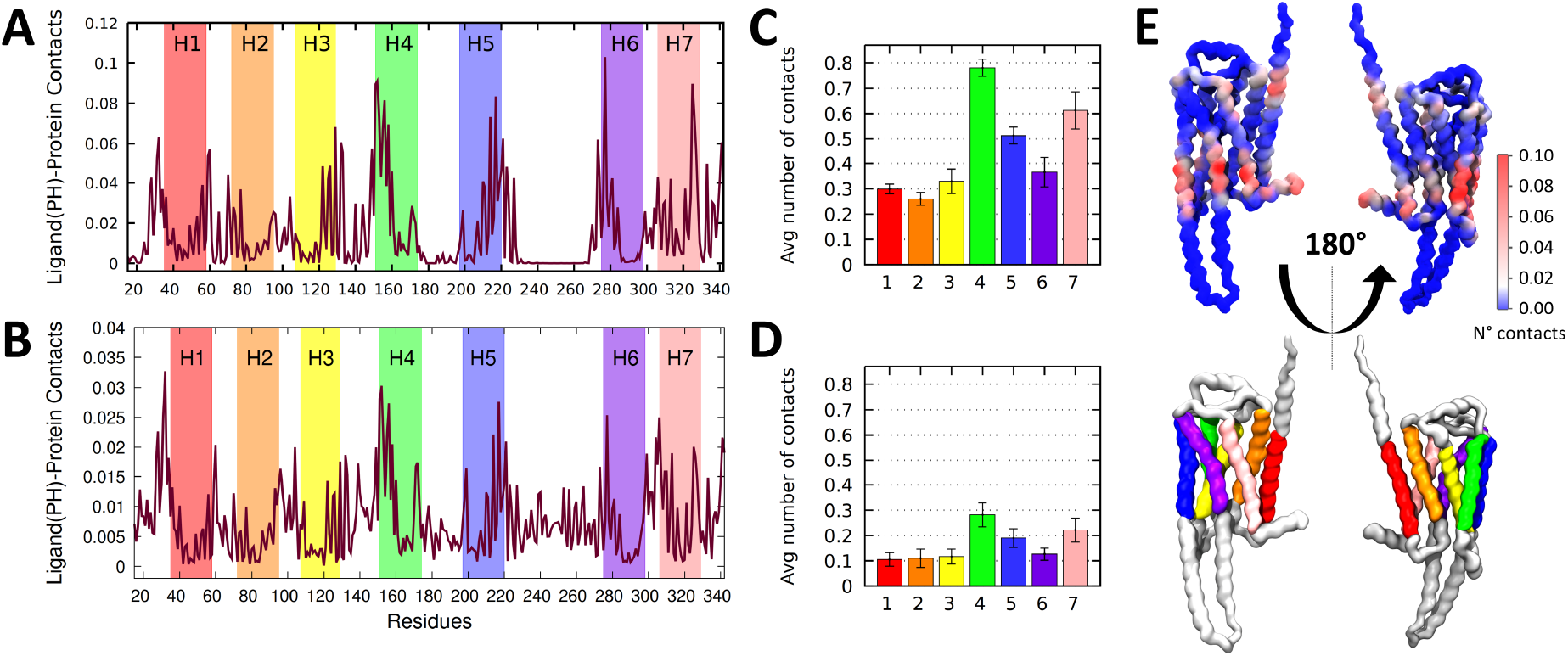
(A) Average number of contacts between the headgroup of SALMT and (B) SALBT with the individual residues of β2AR for the five replicas (25 μs simulation time each). The colored rectangles indicate the seven transmembrane domains (H1-H7) of β2AR. Note the difference in y-range between both plots. (C) Average number of contacts of the transmembrane domains (H1-H7) with SALMT and (D) SALBT headgroup averaged over the five replicas. Error bars show the standard deviation. (E) Front and back view of β2AR colored according to the number of contacts of each residue with the SALMT headgroup (top). The same protein orientations with the transmembrane domains following the color scheme of (A) are depicted below.

To evaluate which helix interacts the most with the drug molecules, the bar graphs in Figure 3C and D show the number of contacts for each transmembrane helix averaged over the five replicas. Both drug molecules show a preference for helix H4, followed by H7 and H5. Minor differences are present for their relative preference of H1, H2 and H3. The overall higher average contact number of SALMT can be attributed to a higher concentration in the membrane due to its higher lipophilicity. This also explains the small number of contacts of SALMT with the extracellular or intracellular loops. The contacts of SALBT are more equally spread over the protein surface (see Figure S15 in the SI).

Figure 3E shows the SALMT-protein contacts on the protein structure. Overall, the parts of the transmembrane helices which are oriented towards the cell interior and in contact with the lower leaflet in the depicted orientation of the protein, show more contacts. This is particularly pronounced for helix H4 (green), where the parts with high contact numbers deeply expand into the membrane. A similar picture emerges for the transmembrane helices of β2AR with SALBT (see Figure S16 in the SI).

We hypothesized that the increased number of *flip-flops* in the presence of β2AR is due to a reduced free energy barrier for membrane crossing of the drug molecules. To quantify the difference between the free energy barriers in the pure membrane and on the β2AR surface, we conducted US calculations. To generate configurations along the reaction coordinate for a pure POPC membrane, the drug molecules were pulled along the membrane normal as explained in detail in Section S1.4 in the SI. Each window was sampled for 2 μs. The resulting potentials of mean force (PMFs) for SALMT (red) and SALBT (blue) are depicted in Figure 4A. As expected, the minima of the PMFs match well the maxima in the density distributions (Figure 1E and F). For both drug molecules, the barrier for *flip-flopping* in the POPC membrane is similar: SALMT has a barrier of 28.0 kJ/mol, and SALBT of 26.5 kJ/mol. In addition, we quantified the difference in lipophilicity between SALMT and SALBT based on their PMFs. In the case of SALBT, the PMF exhibits an energy difference of 4.0 kJ/mol between the minimum in the membrane and water phase, while this difference is 28.5 kJ/mol for SALMT.

**Figure 4:**
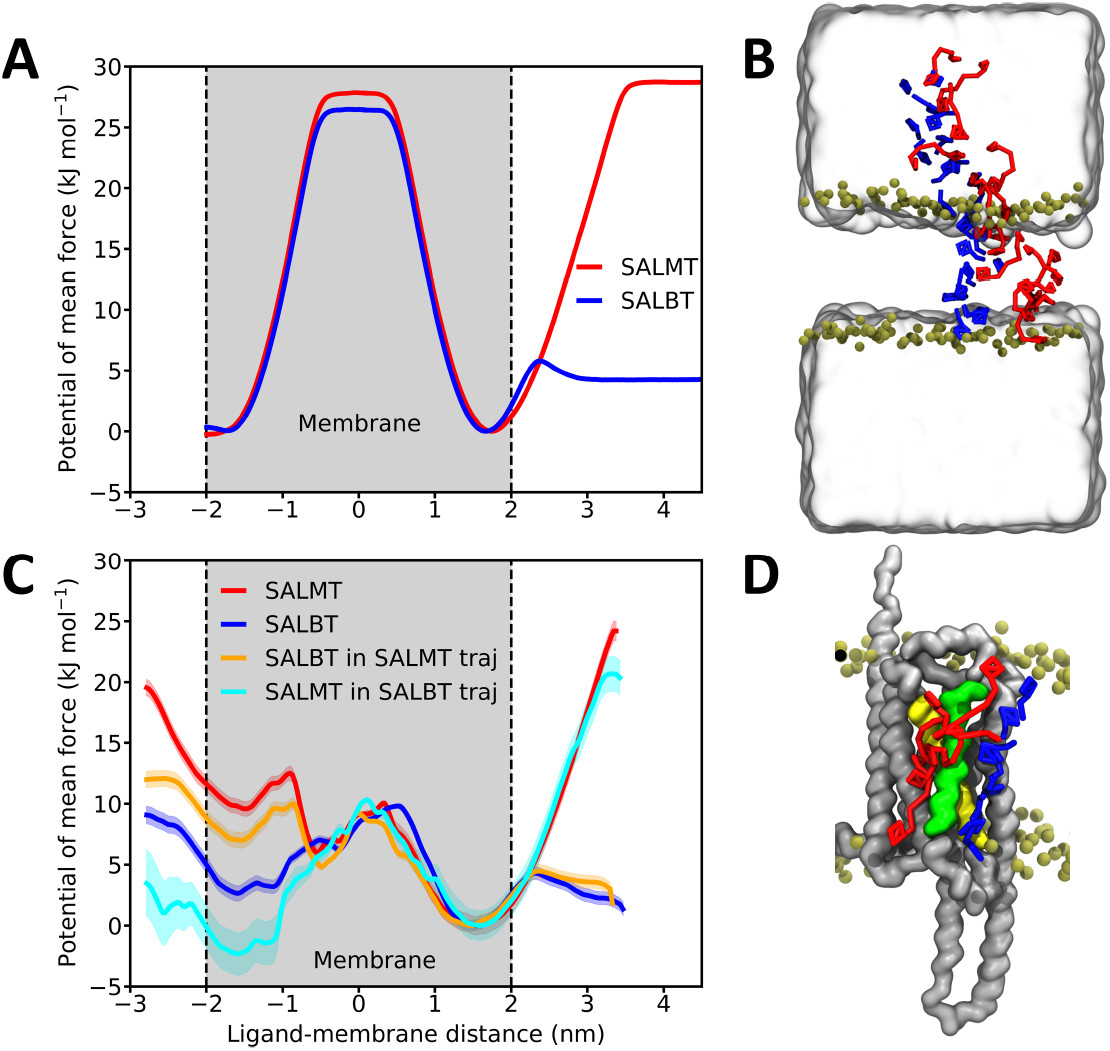
(A) PMFs for SALMT (red) and SALBT (blue) pulled from the water phase towards the centre of a POPC membrane. Error bars calculated with bootstrapping are below the linewidth. (B) Snapshots of the pulling trajectory of SALMT (red) and SALBT (blue). The lipids’ PO4 beads are shown as green spheres indicating the position of the membrane; water is shown as transparent surface. (C) PMFs for SALMT (red) and SALBT (blue) calculated along an unbiased *flip-flop* path on the protein surface. The PMFs for SALBT following the path of SALMT (orange), and for SALMT following the path of SALBT (cyan) are depicted as well. For all PMFs the minimum at positive z-values is set to 0 kJ mol^*−*1^. The shaded areas show the error calculated with bootstrapping. (D) The two different unbiased *flip-flop* pathways with the drug molecules being colored as the respective PMF in (C).

To calculate the PMFs for the *flip-flop* on the protein surface, we selected one of the *flip-flops* along helix H4 from the unbiased trajectories for each drug molecule (Figure 4D) and used the snapshots as starting structures for the US. The SALMT *flip-flop* occurred from the lower leaflet (cytosolic) to the upper one (extracellular); and the SALBT *flip-flop* followed the opposite direction. *Flip-flops* along helix H4 were chosen, because it is the helix with the highest number of contacts to the drug molecules. In addition, we replaced the respective drug molecule by the other one to test, to which extent the difference between the two paths impacts the PMFs.

The resulting PMFs are shown in Figure 4C. First, the energy barrier for all depicted PMFs drops substantially from 28 kJ/mol and 26.5 kJ/mol, respectively, obtained in pure POPC to about 10 kJ/mol on the β2AR surface. Assuming the same frequency factor in the Arrhenius equation, the rate constant would increase by approximately three orders of magnitude. Second, the PMFs are not symmetric anymore in the membrane nor in the water phase, due to the asymmetric protein surface which extends from the membrane into the water phase. The exact barrier height depends on the *flip-flop* pathway, encountering an intermediate metastable state at -0.5 nm for the unbiased path of SALMT (red and orange line). Here, the barrier is asymmetric and thus also depends on the *flip-flop* direction. On the other hand, the PMFs along the unbiased path of SALBT (blue and cyan line) exhibit a more similar barrier from both directions. Besides, SALMT simulated along the SALBT path (cyan line) exhibits a lower energy minimum at the negative x-axis values corresponding to the lower cytosolic leaflet. Contrary to the three other cases, this results in a preferred *flip-flop* direction towards the cytosol from outside the cell. Overall, the resultant PMFs express a stronger dependence on the *flip-flop* path than on the drug molecule.

To investigate if drugs targeting other proteins can use the β2AR surface for membrane permeation, we performed unbiased simulations (10 drug molecules, 5 replicas, 25 μs each; for details see Section S1.1 in the SI) with two kinase inhibitors, baricitinib and dasatinib, for which Martini 3 models were readily available [20]. Baricitinib, a non-receptor tyrosine kinase inhibitor, is used to treat, for instance, rheumatoid arthritis, alopecia, and severe cases of COVID-19 [21, 22]. It is characterised by high solubility in organic solvents and poor solubility in aqueous solution which is associated with its low cell permeability. This can affect the bioavailability and thus its performance as a pharmacological compound requiring higher doses of the drug to achieve its therapeutic effects [23]. Meanwhile, dasatinib, a potent tyrosine kinase inhibitor, finds application in combating various forms of cancer, including leukaemia [24]. It exhibits similar physicochemical properties to those of baricitinib with an even lower solubility in aqueous environments and higher in organic solvents. Therefore, cellular penetration might also be a potential challenge for dasatinib. Density distributions along the membrane normal show the high membrane affinity of both kinase inhibitors (see Figures S24 and S25 in the SI). The structure of these drugs differs considerably from SALMT and SALBT as illustrated in Figure 5A and B. Notably, a key distinction is the absence of the polar head common to SALMT and SALBT, the part that is mainly responsible for interacting with the protein surface during the *flip-flop*.

**Figure 5:**
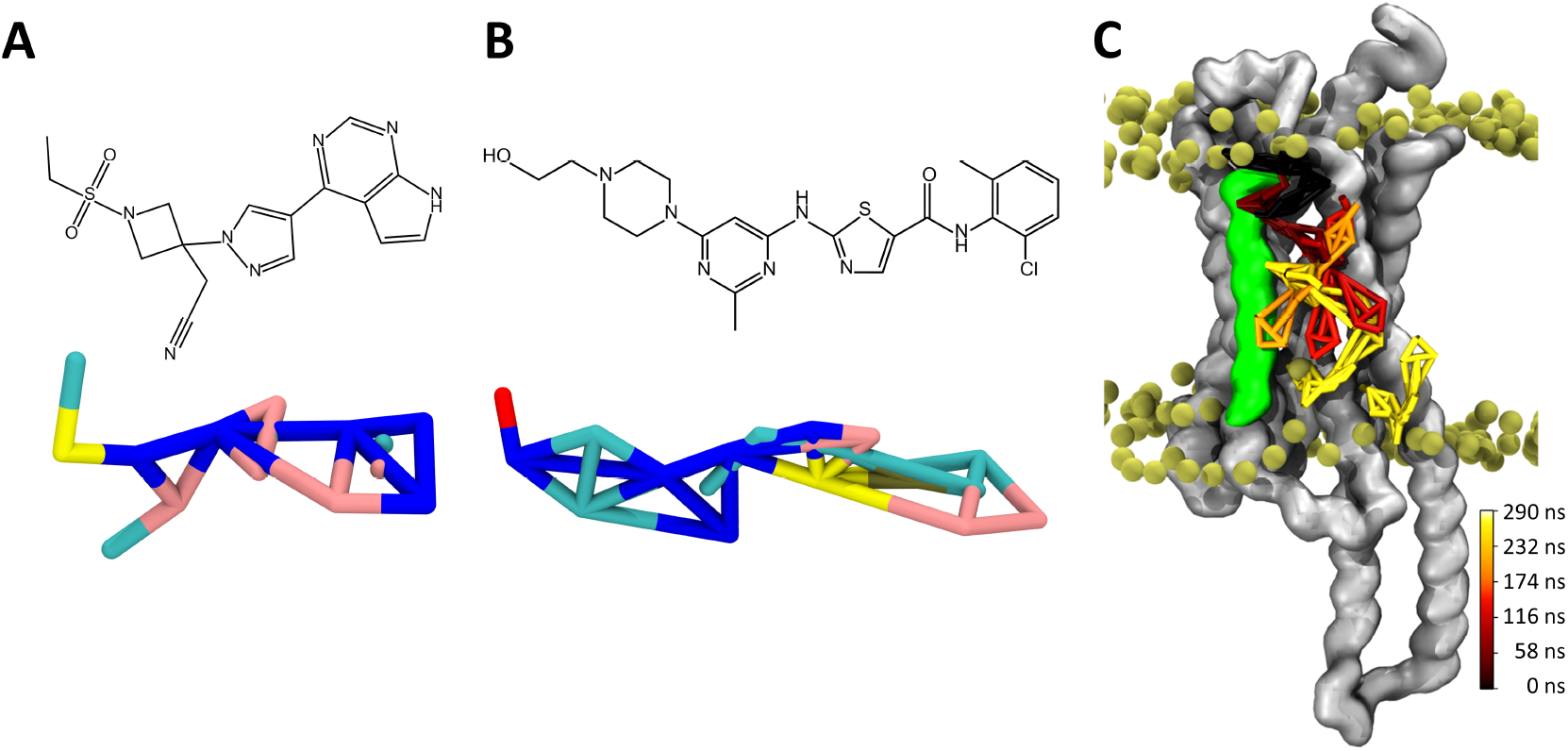
(A) Baricitinib and (B) dasatinib chemical structure (top) and Martini 3 CG model (bottom). (C) Snapshots of a dasatinib *flip-flop* on the protein surface (grey) from the upper (outer) to the lower (cytosolic) leaflet. For the protein, one snapshot is depicted, while for dasatinib several snapshots are shown which are colored from black to yellow representing the evolution in time. The lipids’ PO4 beads are shown as green spheres indicating the position of the membrane.

Our CG simulations revealed that both kinase inhibitors exploit the β2AR surface to change between the membrane leaflets. Over the 125 μs of the five replicas, baricitinib *flip-flopped* two times in each direction. All *flip-flops* occurred on the β2AR surface. In the case of dasatinib, six *flip-flops* happened from outside the membrane towards the cytosol and four in the reverse direction. Of these *flip-flops*, five and three, respectively, occurred on the β2AR surface. Thus, the presence of β2AR also enhances the number of *flip-flops* of the tested kinase inhibitors.

In summary, our results suggest a previously unknown role of β2-adrenergic receptor (β2AR), a class A GPCR, in facilitating the membrane permeation of drug molecules. Our CG MD simulations show that β2AR facilitates membrane permeation of its agonists salbutamol and salmeterol. PMFs for their membrane permeation show a reduction of the energy barrier by approximately 60% from more than 25 kJ/mol to about 10 kJ/mol. We also observed this effect for two kinase inhibitors whose targets reside in the cytosol. Interestingly, GPCRs are not only present in the plasma membrane, they are also found in the blood-brain barrier [25]. This suggests that they could also potentially facilitate drug permeation there [26], making this an exciting new potential design pathway for drugs needing to traverse the plasma membrane or the blood-brain barrier. Our in silico results are supported by the experimentally known scramblase activity of several GPCRs [7, 9].

Moreover, our simulations revealed that the β2AR agonist salmeterol interacts preferably with the protein part embedded in the cytosolic leaflet. Besides salmeterol’s high lipophilicity, this might additionally contribute to the slow-acting behavior of salmeterol. Because the binding pocket is located in the protein part embedded in the extracellular leaflet, salmeterol has to perform a *flip-flop* to reach it.

## Computational Methods

All simulations were performed with the program package GROMACS (version 2020.4)[27]. The coarsegrained simulations were performed with the Martini 3 force field [17]. The all-atom simulations required for the parametrization of the CG models were performed with the CHARMM36 force field [28] in the case of the protein and the OPLS-AA force field [29] in the case of the drug molecule. The PMFs were calculated using US. The starting structures for each window were taken from unbiased simulations in case of the *flip-flop* events on the protein surface or generated by pulling simulations. The simulations were analyzed using GROMACS tools and in-house custom Python scripts. Details of the simulation settings, the parametrization of the CG models, as well as the analysis are provided in the SI.

## Acknowledgements

We thank Saara Lautala for critically reading the manuscript. We acknowledge the Alfons und Gertrud Kassel Foundation, the Dr. Rolf M. Schwiete Foundation, and the Center for Multiscale Modeling in Life Sciences (CMMS) sponsored by the Hessian Ministry of Science and Art for funding and the Center for Scientific Computing at Goethe University Frankfurt for access to Goethe-HLR and FUCHS.

## Supporting Information

The Supporting Information is available free of charge at https://pubs.acs.org/.

1. Computational methods (computational and simulation details, drug molecule and protein parametrization, analysis, free energy calculations)
2. Parametrization of drug molecules models
3. Itp files
4. Parametrization of the protein β2AR
5. Scheme of the counting *flip-flop* script threshold
6. SALMT behavior in the membrane
7. Drug molecule behavior in the membrane with β2AR (Distance of the drug molecules to the middle of the membrane)
8. Flat-bottom potential for SALBT simulations (number of flip-flops, density plots, and distance plots of the ligands to the middle of the membrane)
9. Flip-flop Localization on β2AR
10. US convergence tests and histograms
11. Dasatinib and baricitinib density plots

