## Supplemental Material for "GPCR surface creates a favorable pathway for membrane permeation of drug molecules"

March 15, 2024

**2 Parametrization of drug molecule models** **6**

**3 Itp files** **10**

**4 Parametrization of the protein: β2AR** **12**

**5 Scheme of the counting *flip-flop* script thresholds** **12**

**6 SALMT behavior in the membrane** **13**

**7 Drug molecule behavior in the membrane with β2AR** **14**
- 7.1 Distance of the drug molecules to the middle of the membrane . . . 15

**8 Flat-bottom potential for SALBT simulations: Number of flip-flops, density plot, and distance of the ligands to the middle of the membrane** **16**

**9 Flip-flop localization on β2AR** **20**

**10 US convergence tests and histograms** **22**

**11 Dasatinib and baricitinib density plots** **25**

### 1 Computational methods

#### 1.1 Computational and simulation details

The simulations presented in this work were performed using the program package GROMACS (version 2020.4) [1]. The Martini 3 force field [2] was employed for all coarse-grained (CG) simulations. For the all-atom simulations used as reference for the parametrization of the CG models, the CHARMM36 force field [3] was applied for the protein simulations and the OPLS-AA [4] for the drug molecule simulations. All systems were neutralized and solvated in 0.15 M NaCl solution, except for the simulations for the drug molecule parametrization where no ions were included. CG systems were set up using the script *insane.py* [5]; atomistic systems were set up using CHARMM-GUI [6, 7, 8]. In the CG membrane systems without proteins, 100 1-palmitoyl-2-oleoyl-sn-glycero-3-phosphocholine (POPC) lipids composed each of the two leaflets that together with the 10 drug molecules (salmeterol, salbutamol, dasatinib, baricitinb) were solvated with 2,700 water beads, 30 $Na^+$ beads and 30 $Cl^-$ beads. The total box dimensions were approximately 8 nm × 8 nm × 9 nm. The CG systems when β2-adrenergic receptor (β2AR) was included consisted of 200 POPC lipids in each leaflet, 10 drug molecules (salmeterol, salbutamol, dasatinib, baricitinb), 11,000 water beads, 120 $Na^+$ beads and 120 $Cl^-$ beads. The total box dimensions were approximately 12 nm × 12 nm × 13 nm. For the drug molecule parametrization the system consisted of one drug molecule and 900 water beads in a box of 5 nm × 5 nm × 5 nm and for the protein parametrization the system increased to 12 nm × 12 nm × 16 nm and consisted of 200 POPC lipids in each leaflet, 14,000 water beads, 150 $Na^+$ beads and 150 $Cl^-$ beads. The all-atom systems used for reference comprised one drug molecule and 1,700 water molecules in a box size of 4 nm × 4 nm × 4 nm; the protein systems additionally included 150 POPC lipids in total, 44 $Na^+$ ions, 47 $Cl^-$ ions and 17,000 water molecules.

For all Martini CG simulations, an initial steepest descent minimization of 2000 steps was performed. Afterwards, an equilibration was carried out in the NPT ensemble employing the v-rescale thermostat with a reference temperature of $T_{\text{ref}} = 298$ K ($\tau_T = 1.0$ ps), and the Berendsen barostat with a reference pressure of $p_{\text{ref}} = 1$ bar ($\tau_p = 5$ ps). A timestep of $\Delta t = 10$ fs was used for the equilibration. In the systems without protein, the equilibration was 500 ps long; in the systems containing proteins, it was 2.5 ns long.

Most of the simulation parameters were kept for the production phase, except for the barostat, which was set to Parrinello-Rahman with an increased coupling constant of $\tau_p = 12$ ps. Van der Waals and Coulomb interactions were always treated with a cutoff scheme (1.1 nm) and Coulomb interactions were treated with reaction-field. The timestep was increased to $\Delta t = 20$ fs in the 25 µs long production runs for all systems. To ensure a proper stability of the parametrized molecules, a timestep of $\Delta t = 20$ fs and $\Delta t = 30$ fs were used for the equilibration and production runs of the drug molecule parametrization, respectively. The length of the production runs was 75 ns.

In the atomistic reference simulation for the drug molecule parametrization, a steepest descent minimization of 500 steps was run. A 250 ps equilibration with a timestep of $\Delta t = 1$ fs was performed in an NPT ensemble, using the Berendsen thermostat and barostat with a coupling constant of $\tau_p = 0.5$ ps to keep the same temperature and pressure as in the CG simulations ($T_{\text{ref}} = 298$ K, $p_{\text{ref}} = 1$ bar). In the production runs, the timestep was raised to $\Delta t = 2$ fs in the 50 ns production runs and the thermostat was switched to Nosé–Hoover and the barostat to Parrinello-Rahman (coupling constants $\tau_T = 1$ ps and $\tau_p = 5$ ps). Van der Waals interactions were treated with a cutoff scheme (1.4 nm) and

Coulomb interactions were calculated using reaction field with a cutoff scheme (1.4 nm) according to the recommended settings with the OPLS-AA force field. In the atomistic simulation of the protein, the minimization phase was increased to 5000 steps, and dividing the equilibration into six steps to slowly release the position restraints established in all the protein and membrane and increasing the timestep from $\Delta t = 1$ fs to $\Delta t = 2$ fs. The first three equilibration steps were carried out in an NVT ensemble for 125 ps each applying the Berendsen thermostat ($\tau_T = 1$ ps), and the last three in an NPT ensemble for 500 ps each applying a Berendsen thermostat ($\tau_T = 1$ ps) and a Berendsen barostat with a coupling constant of $\tau_p = 5$ ps to keep $T = 298$ K and $p = 1$ bar. For the 1 μs production runs, the $\Delta t = 2$ fs timestep was maintained, and the temperature and pressure were kept with the Nosé-Hoover barostat and the Parrinello-Rahman thermostat ($\tau_p = 5$ ps). Van der Waals interactions were treated with a cutoff scheme (1.2 nm); Coulomb interactions were calculated using PME (1.2 nm) according to the recommended settings for the CHARMM force field.

#### 1.2 Drug molecule and protein parametrization

For the parametrization of the drug molecules salmeterol (SALMT) and salbutamol (SALBT), the detailed procedure is described in the chapter by Alessandri et al.[9]. Firstly, all-atom simulations of the drug molecule in water solvent and in hexadecane were performed, obtaining the drug molecule parameters from the Parameter Generator for Organic Ligands (LigParGen server [10, 11, 12]). From these simulations, a CG trajectory is generated mapping the all-atom structures to CG resolution. From this mapped trajectory we extracted the bond, angle, and dihedral angle distributions. They served to build an initial CG model which was subsequently refined. Then, the volume and solvent partitioning of the drug molecule model were compared to the volume of the all-atom model and the experimental *log P* data, respectively. The solvent-accessible surface areas (SASAs) were calculated with the GROMACS tool *gmx sasa*. The octanol-water partitioning was estimated from the individual solvation free energies in both solvents. The latter were calculated by performing a thermodynamic integration using GROMACS, which was analyzed with the alchemlyb Python package [13]. The behavior of the final drug molecule models in presence of a POPC membrane was tested in five simulation replicas of 25 μs each.

For the parametrization of the β2AR protein, a different procedure was implemented [9]. The atomistic protein structure was obtained from the AlphaFold Protein Structure Database [14]. The disordered regions at the termini (1MET-14PRO and 343ARG-413LEU) were removed since these regions are not expected to play a relevant role in the drug molecule *flip-flop*. Then, using the Martini tools *martinize2* [15] and *create_goVirt.py* [?], its CG structure and topology files were generated. As recommended for Martini 3 proteins, the side chain correction flag *-scifix* was specified using *martinize2*. To maintain the secondary and tertiary structure elements, the GōMartini model was used as the structure bias model [16, 17]. The added Gō-bonds were described by a Lennard-Jones potential between the backbone beads of residues with a natural contact and a distance of 0.3 to 1.1 nm in the CG reference structure of β2AR. Using virtual interaction sites, the dissociation energy of the interactions was fixed at ε= 10.0 kJ/mol. The total number of Gō-bonds added was 615.

#### 1.3 Analysis

**Flexibility comparison.** To compare the flexibility between atomistic and CG protein models (Figure S3, the trajectories were iteratively fitted to an increasing number of $C_\alpha$ atoms and backbone beads, respectively, which were the most rigid ones in the structure. Initially, the trajectory was processed in order to keep the protein whole and center it, and the RMSF was calculated using the GROMACS tool *gmx rmsf* after fitting the rotation and translation of all the $C_\alpha$ atoms of β2AR, 328. Subsequently, the $C_\alpha$s with an RMSF below a threshold of 1 Å were utilized for fitting the trajectory in the next iteration. This iterative procedure was repeated until the list of rigid $C_\alpha$ atoms converged. For the final RMSF profiles, 135 $C_\alpha$ atoms (out of the total 328) were considered the most rigid ones and thus, used for fitting. Analogously, the RMSF of the GōMartini protein model was computed using the backbone beads for fitting. Out of the total number of 328 backbone beads, 184 were used for fitting.

**Counting *flip-flops*.** The computation of the number of *flip-flops* was performed using a combination of GROMACS tools and custom Python scripts. First, *gmx distance* was employed to determine the distance between the center of mass (COM) of the terminal beads of the lipid tails (C4A and C4B) and the COM of the drug molecule. In the following steps, only the component in the direction of the membrane normal (the z-component here) of the resulting distance vector is considered. Subsequently, the z-components were classified into four different areas: outside the membrane, in the membrane middle, in the upper leaflet, or in the lower one. The area outside the membrane was defined as $z > 2.5$ nm and $z < -2.5$ nm, while the middle of the membrane was defined as $1.0 \text{ nm} > z > -1.0$ nm. The upper and lower leaflets were the remaining regions in positive and negative ranges, respectively, as depicted schematically in Figure S4 for a SALMT (left) and a SALBT trajectory (right). The resulting status of the drug molecule is stored and evaluated along with the next z-value. A change in leaflet is only counted as *flip-flop* if the drug molecule status changes from one leaflet to the opposite via the membrane middle but not via the region outside the membrane. When the protein is also present in the simulations, the analysis is slightly augmented in order to differentiate *flip-flops* on the protein surface from the ones in the bilayer. For that, the GROMACS tool *gmx distance* is required to compute the 3D-vector between the COM of the protein and the COM of the drug molecule. Every time that a *flip-flop* is identified, the distance to the protein, i.e. the norm of the vector between the COMs, is evaluated. When this distance is lower than 3.8 nm, which is approximately 1 nm larger than the protein radius (the maximum distance at which intermolecular interactions are computed), the *flip-flop* is considered to take place on the protein surface. Both scripts were verified by visual inspection of the computed graphs of the z-component of the drug molecule-lipid tail end distance as the examples in figures S8 and S9.

**Drug molecule-protein contacts.** The analysis of the number of contacts of the drug molecules and the protein throughout the simulation was performed employing custom shell and Python scripts in combination with GROMACS tools. The part of the drug molecule considered for this analysis is the polar head of SALMT and SALBT, which is identical in both molecules. First, the number of contacts between the polar head and each protein residue is calculated for a distance cutoff of 0.7 nm using *gmx select*. Then, the contact fraction is calculated by dividing the number of contacts per residue by the number of frames for each replica, which was further averaged over the five replicas. Finally, we also computed the mean value for each transmembrane domain to analyze in which parts of the protein most of the drug molecule contacts occur.

**Figures.** Figures were prepared with the Python library Matplotlib [18], Gnuplot[19], and the program package Visual Molecular Dynamics (VMD) [20].

#### 1.4 Free energy calculations

To estimate the free energy of the *flip-flop* process in the pure membrane and on the protein surface, we calculated potentials of mean force (PMFs) using umbrella sampling (US). In case of the pure membrane, we firstly pulled the drug molecules from the water phase to the middle of the membrane in order to generate the initial conformations for the US windows [$z$-direction; force constant = 1000 kJ/(mol nm$^2$); pulling rate = −0.001 nm/ps]. The reaction coordinate was defined as the distance along the z-axis between the COM of the five beads composing the head of the drug molecules and the COM of the group composed of the terminal beads in the lipid tails (C4A and C4B). For the US, 59 windows were run for SALMT from 5.8 to 0.0 nm, and 57 windows for SALBT, from 5.6 to 0.0 nm. In both cases, the windows had a distance of 0.1 nm and were sampled for 2 µs. The distance between the drug molecule and the lipid tails was restrained with a harmonic potential using a force constant of 1000 kJ/(mol nm$^2$). For sampling the entire membrane, a second pulling with the same parameters was performed in the opposite direction starting in the middle of the membrane until the phosphate and choline beads (PO4, NC3) of the other bilayer. 21 windows were added for each of the drug molecules from 0.0 to -2.0 nm. This allows us to judge the convergence of the PMF in both leaflets as well as to compare the PMFs with the systems including β2AR.

For the US simulations including the protein, snapshots were extracted along a representative *flip-flop* path obtained in the unbiased simulations and used as initial configurations of the umbrella windows. In addition, to complete the sampling of the membrane and the water phase along the z-direction and for a better comparison with the US of the systems without the protein, two additional pullings of the drug molecules were performed. For that, the drug molecules were pulled from the last snapshot at each of the membrane edges to penetrate the water phase in both directions. The reaction coordinate for all US simulations along the β2AR surface was defined as the difference between the COM of the polar heads of the drug molecules and the COM of a group of protein beads located right underneath the membrane (62GLU, 135ILE, 136THR, 137SER, 140LYS, 267LYS, 269HIS). The partial density of these beads with respect to POPC and the lower leaflet beads is depicted in Figure S23. A second bias force was implemented to prevent the drug molecules from diffusing away from the β2AR surface. This bias force restrained the distance in x- and y-direction between the COM of the drug molecules' polar heads and the COM of the residues in the transmembrane region around the *flip-flop* path: H3 (residue 107-129), H4 (residue 151-274) and H5 (residue 197-220). For SALMT, 80 umbrella windows were run using snapshots taken from the unbiased *flip-flop*, 18 more windows were added from the pulling at the lower part and 32 at the upper part. In the case of SALBT, 77 windows were run using from snapshots from the unbiased *flip-flop*, 15 were added at the lower part and 29 at the upper part. Each of the windows was run for 2 µs. The z-distance between the windows was around 0.05 nm, resulting in an overall sampling from $(-3)-3.5$ nm considering zero as the bilayer center. Two more US calculations were run with the same amount of windows but substituting SALMT for SALBT in each of the initial snapshots of the US and vice versa. In this way, the PMFs were calculated in an identical manner for both drug molecules when following their unbiased *flip-flop* paths as well as the one from the other drug molecule.

All PMFs were calculated using the *gmx wham* tool. The initial 200 ns of each umbrella window were discarded. Error estimation was done using a bootstrap analysis with 100 bootstraps [21]. The convergence behavior of the PMFs is shown in Section 9, Figures S17−S22). Overall, 316 µs of US simulations in the pure POPC membrane and 502 µs on the protein surface were performed for both drug molecules in total.

#### 2 Parametrization of drug molecule models

In Martini 3 force field, aromatic structures are best described by tiny (T) beads. For pairs of non-substituted aromatic C atoms, the recommended bead type is TC5. Thus, beads R3, R4, R9, R10, and R11 of the two aromatic moieties in SALMT are defined as TC5 beads. The two hydroxyl groups mapped into the beads O1 and O2, are represented by a SP1 and TN6 bead, respectively. The SP1 bead represents the higher polarity of the aliphatic hydroxyl group and has a small (S) size due to the mapping of three non-hydrogen atoms to one CG bead (3-to-1 mapping). The aromatic hydroxyl group in O2 is slightly less polar and thus represented by a TN6 bead [22].

Linear alkanes are typically represented by C1 beads, thus, bead R8 is of SC1 type. The small (S) bead was chosen due to the 3-to-1 mapping. Although bead R6 also describes a linear alkane moiety, it was defined as a regular C2 bead, because of its proximity to the amine group which renders it slightly more polar.

The ether is described by a regular N3a bead with a 4-to-1 mapping. The most challenging bead type assignment was the one of bead N5, because it includes an alcohol group and a secondary amine. Therefore, a bead type more polar than the one defined for an alcohol (P1) had to be chosen. After testing the free energy of transfer from hydrated octanol to water for the beads P3-P6 (see Table S1), an SP5 bead type was chosen as it reproduced well enough the energy without deviating much from the recommended bead type involving those chemical groups.

The bonded terms of the aromatic rings were described by constraints, while the flexible linker was described by harmonic bonds. For the functionalized aromatic ring (O1-R4), a hinge model was used [22, 23]. To maintain the molecule in a linear conformation, angles were introduced between every three beads, applying a subtle force constant to also account for the bent form. Figure S1 depicts the probability distributions for the bonded terms of the atomistic and the CG model, which are in good agreement. To ensure an accurate representation of the molecular volume, we also calculated the SASA values. The SASA of the CG model of $9.19 \pm 0.38$ nm$^2$ reproduces well the atomistic value of $9.27 \pm 0.60$ nm$^2$.

The logarithm of the experimental hydrated octanol/water partition coefficient found in the literature for SALMT ranges from 3.61 [24] to 4.2 [25]. This coefficient corresponds to a free energy of transfer from hydrated octanol to water of $\Delta G^{\mathrm{exp}}_{\mathrm{oct/w}} = 20.73 - 24.40$ kJ/mol according to $logP = \Delta G_{\mathrm{oct/w}}/ln(10)RT$ [25]. For the optimized Martini 3 model of this drug molecule, SALMT, the free energy of transfer is $\Delta G^{\mathrm{CG}}_{\mathrm{oct/w}} = 25.54 \pm 0.22$ kJ/mol which is in good agreement with the experimental value. The Martini 3 model of SALBT was generated based on the SALMT model (see Figure 1C). The probability distributions for the bonded terms (see Figure S2) and the SASA value (CG: $5.66 \pm 0.12$ nm$^2$; AA: $5.33 \pm 0.14$ nm$^2$) compare well to the atomistic reference. Its calculated free energy of transfer of $\Delta G^{\mathrm{CG}}_{\mathrm{oct/w}} = -3.61 \pm 0.15$ kJ/mol is comparable with the range of experimental values found, $logP = 0.3 - 0.6$ ($\Delta G^{\mathrm{exp}}_{\mathrm{oct/w}} = 1.72 - 3.68$ kJ/mol)[25, 26]. While certain bead types might yield slightly more accurate free energies of transfer, we maintained consistency in bead selection for both drug molecules, especially given the structural homogeneity up to the R6 bead. Given the slightly wider range of experimental values encountered, the bead types commonly representing these groups of atoms in Martini 3 were preferred for SALMT and SALBT. Concerning SALMT, an SP6 bead type for bead N5 is overly polar for a secondary alcohol and a secondary amine, therefore, SP5 was chosen. In the case of SALBT, opting for a smaller bead type for R6 better captured the volume, and a C2 type is usually applied for modelling branched alkanes.

Table S1: Free energy of transfer (hydrated octanol/water) for the different tested bead types for the N5 bead of SALMT and the R6 bead of SALBT, respectively, along with the range of experimental values.

| | N5 Bead Type | R6 Bead Type | $\Delta G_{\mathrm{oct/w}}$ (kJ/mol) | $\Delta G^{\mathrm{exp}}_{\mathrm{oct/w}}$ (kJ/mol) |
|---|---|---|---|---|
| SALMT | SP3 | RC2 | 28.70 ±0.21 | 20.73 − 24.40 kJ/mol |
| | SP4 | RC2 | 26.32 ±0.21 | |
| | SP5 | RC2 | 25.54 ±0.22 | |
| | SP6 | RC2 | 24.86 ±0.20 | |
| SALBT | SP5 | RC1 | 3.55 ±0.14 | 1.72 − 3.68 kJ/mol |
| | SP5 | RC2 | 0.18 ±0.14 | |
| | SP5 | SC1 | −0.47 ±0.14 | |
| | SP5 | SC2 | −3.61 ±0.15 | |

### Parametrization of the drug molecules: bonded terms of the SALMT model

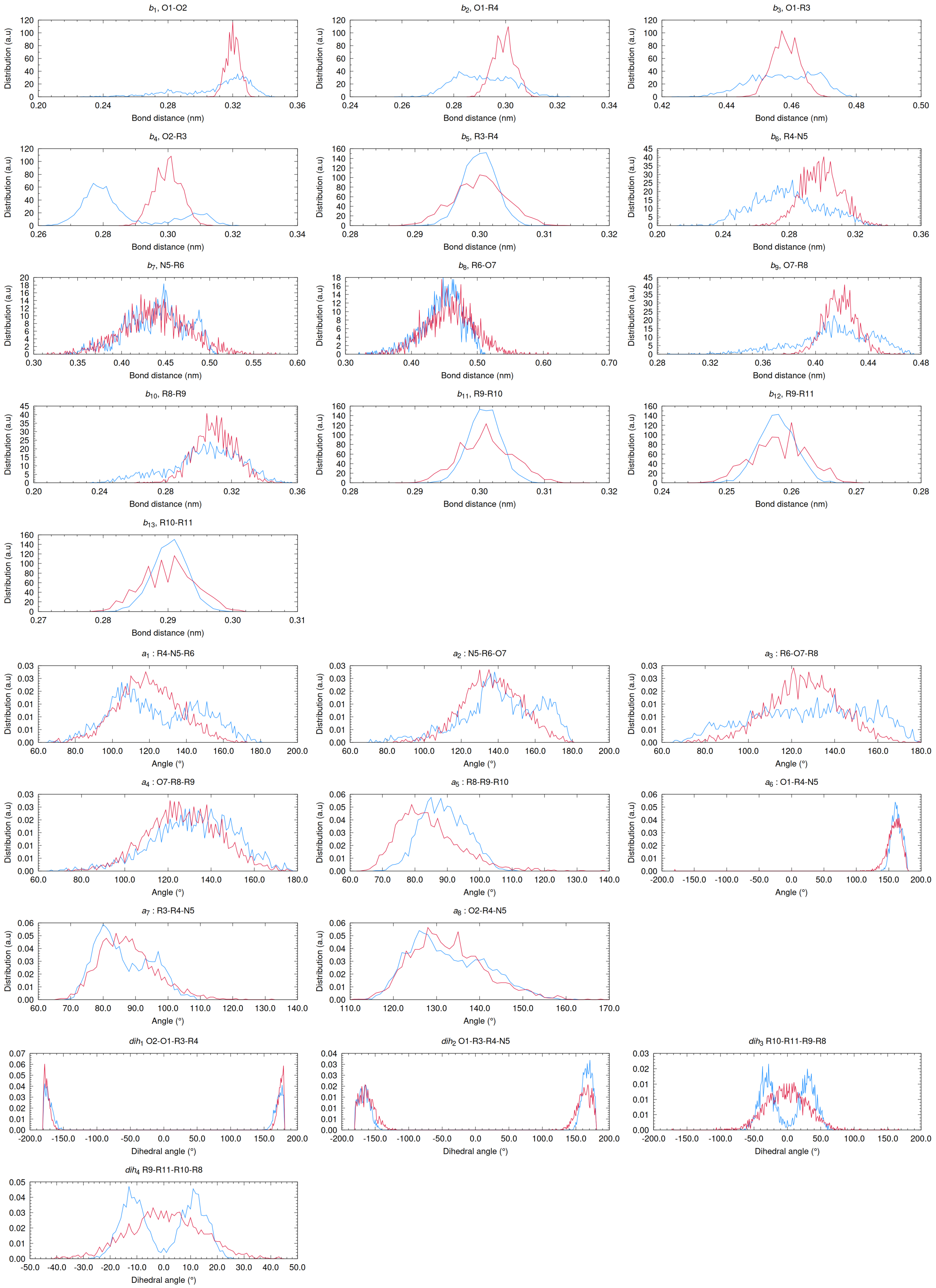

Figure S1: Bond distances (b1-b13), angles (a1-a8) and dihedrals (dh1-dh4) distributions of relevance for SALMT's model. All-atom distributions are displayed in blue and Martini coarse-grained distributions in red.

#### Parametrization of the drug molecules: bonded terms of the SALBT model

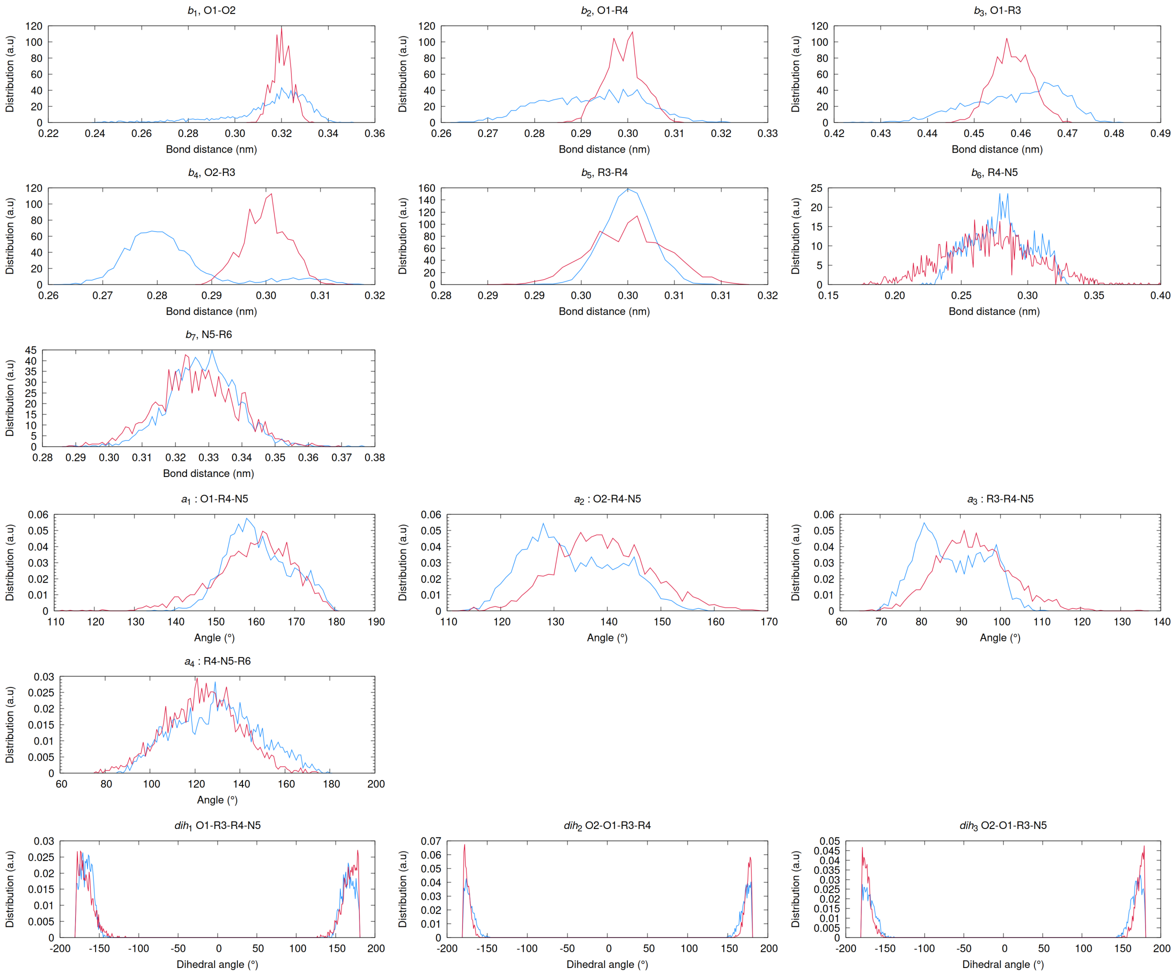

Figure S2: Bond distances (b1-b7), angles (a1-a4) and dihedrals (dh1-dh3) of relevance for SALBT parametrization. All-atom distributions are displayed in blue and Martini coarse-grained distributions in red.

### 3 Itp files

#### SALMT itp file

```
[ moleculetype ]
; molname nrexcl
SALMT 1
[ atoms ]
; nr type resnr residue atom cgnr charge mass
1 SP1 0 SALMT O1 1 0 31
2 TN6 0 SALMT O2 2 0 29
3 TC5 0 SALMT R3 3 0 26
4 TC5 0 SALMT R4 4 0 25
5 SP5 0 SALMT N5 5 0 59
6 C2 0 SALMT R6 6 0 56
7 N3a 0 SALMT O7 7 0 58
8 SC1 0 SALMT R8 8 0 42
9 TC5 0 SALMT R9 9 0 25
10 TC5 0 SALMT R10 10 0 26
11 TC5 0 SALMT R11 11 0 26

[ bonds]
; i j funct length
4 5 1 0.300 20000
5 6 1 0.450 2000
6 7 1 0.470 2000
7 8 1 0.420 20000
8 9 1 0.310 20000
#ifndef FLEXIBLE
[ constraints]
#endif
; i j funct length
1 2 1 0.320 1000000
1 4 1 0.299 1000000
1 3 1 0.458 1000000
2 3 1 0.300 1000000
3 4 1 0.300 1000000
9 10 1 0.301 1000000
9 11 1 0.258 1000000
10 11 1 0.290 1000000

[ angles ]
; ai aj ak funct angle force c.
4 5 6 1 125.000 25 ;
5 6 7 1 145.000 25 ;
6 7 8 1 135.000 25 ;
7 8 9 1 135.000 25 ;
8 9 10 1 70.000 25 ;
1 4 5 1 140.000 25 ;
3 4 5 1 70.000 25 ;
2 4 5 1 120.000 25 ;

[ dihedrals ]
; improper
; i j k l funct ref.angle force_k
2 1 3 4 2 180.000 200
2 1 3 5 2 180.000 100
```

```
8 9 10 11 2 180.000 25

[ exclusions ]
2 4
```

#### SALBT itp file

```
[ moleculetype ]
; molname nrexcl
SALBT 1
[ atoms ]
; nr type resnr residue atom cgnr charge mass
1 SP1 0 SALBT O1 1 0 31
2 TN6 0 SALBT O2 2 0 29
3 TC5 0 SALBT R3 3 0 26
4 TC5 0 SALBT R4 4 0 25
5 SP5 0 SALBT N5 5 0 59
6 SC2 0 SALBT R6 6 0 57

[ bonds ]
; i j funct length
4 5 1 0.280 2000
5 6 1 0.328 20000
#ifndef FLEXIBLE
[ constraints ]
#endif
; i j funct length
1 2 1 0.320
1 4 1 0.299
1 3 1 0.458
2 3 1 0.300
3 4 1 0.300

[ angles ]
; ai aj ak funct angle force c.
1 4 5 1 160.000 25
2 4 5 1 133.000 25
3 4 5 1 88.000 25
4 5 6 1 130.000 25

[ dihedrals ]
; improper
; i j k l funct ref.angle force_k
2 1 3 4 2 180.000 200
2 1 3 5 2 180.000 100

[ exclusions ]
2 4
```

#### 4 Parametrization of the protein: β2AR

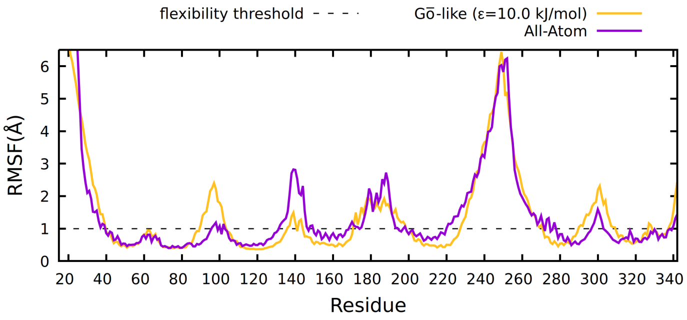

Figure S3: Flexibility comparison of the employed Gō-like model and the all-atom simulation along 1 and 1.5 µs simulations respectively. The first 200 ns of the all-atom simulation were excluded because the system was still equilibrating.

#### 5 Scheme of the counting *flip-flop* script thresholds

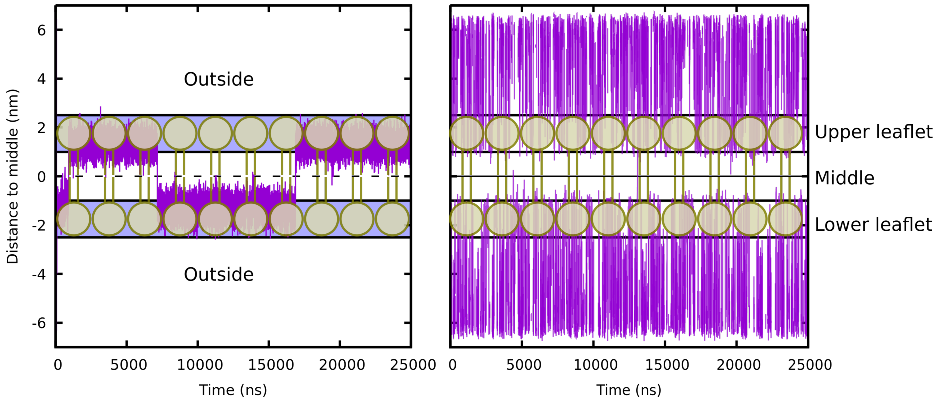

Figure S4: Distance of the center of mass (COM) of SALMT (left) and SALBT (right) drug molecules to the COM of the beads in the lipid tails along one of the 25 µs simulation replica. Displaying a scheme of where the lipids are located and the boundaries of the four regions for counting *flipflops*.

#### 6 SALMT behavior in the membrane

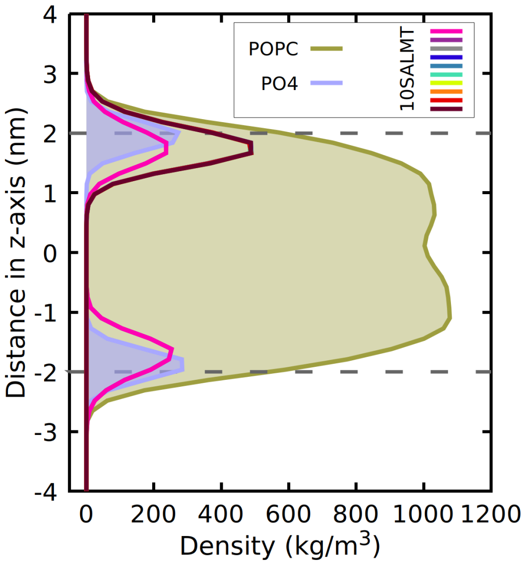

Figure S5: Density distribution of ten SALMT molecules (only considering the polar head of the molecules; beads O1 O2 R3 R4 N5, for comparison with SALBT), the POPC membrane, and the lipids' PO4 beads depicted along the membrane normal (z-axis) from a 25 µs simulation trajectory. The intracellular region corresponds to negative z-values; the extracellular region to positive z-values. The densities of the SALMT molecules are scaled by a factor of 100.

### 7 Drug molecule behavior in the membrane with β2AR

#### SALMT

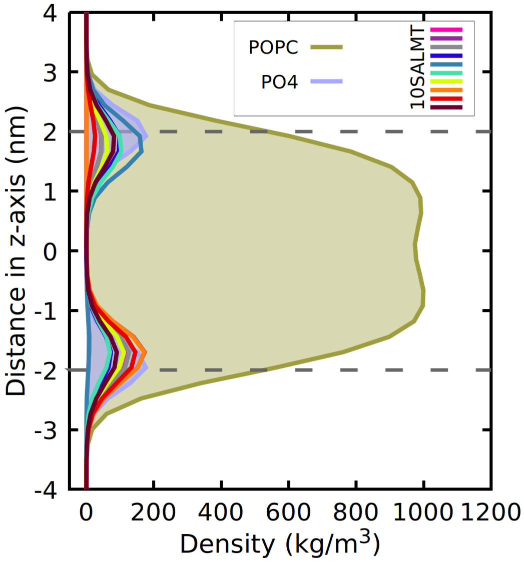

Figure S6: Density distribution of ten SALMT molecules (only considering the polar head of the molecules; beads O1 O2 R3 R4 N5, for comparison with SALBT), the POPC membrane, and the lipids' PO4 beads depicted along the membrane normal (z-axis) from a 25 µs simulation trajectory. The intracellular region corresponds to negative z-values; the extracellular region to positive z-values. The densities of the SALMT molecules are scaled by a factor of 100.

#### SALBT

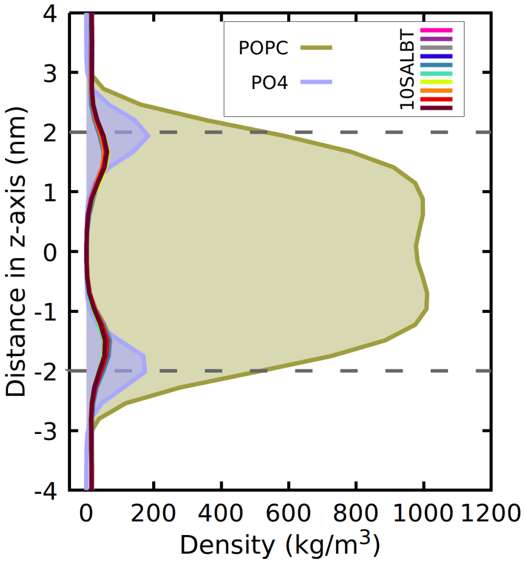

Figure S7: Density distribution of ten SALBT molecules, the POPC membrane, and the lipids' PO4 beads depicted along the membrane normal (z-axis) from a 25 µs simulation trajectory. The intracellular region corresponds to negative z-values; the extracellular region to positive z-values. The densities of the SALBT molecules are scaled by a factor of 100.

#### 7.1 Distance of the drug molecules to the middle of the membrane

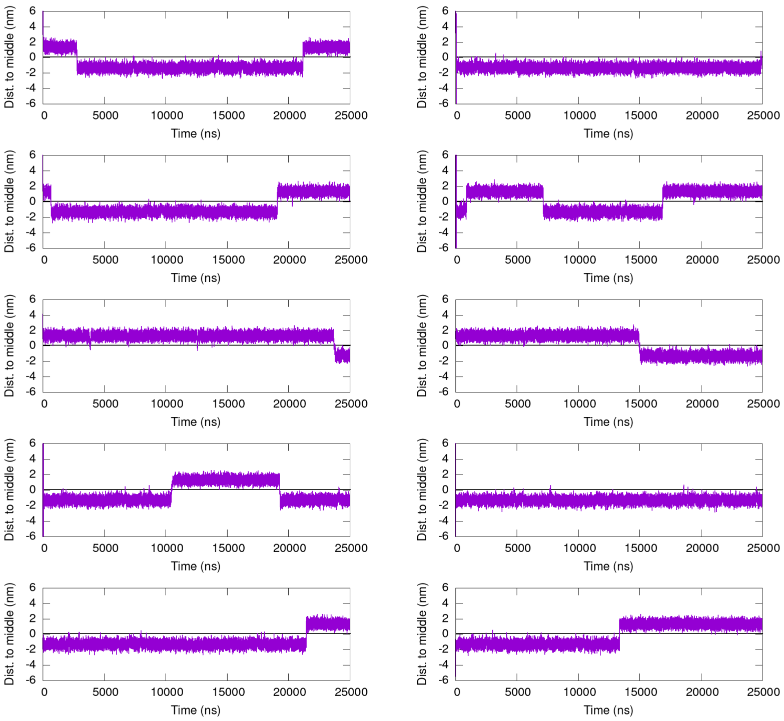

Figure S8: Distance of the COM of each of the 10 SALMT drug molecules to the middle of the membrane along the simulation for one of the replicas in the presence of β2AR. *Flip-flops* can be observed when the distance changes from positive to negative values or vice versa.

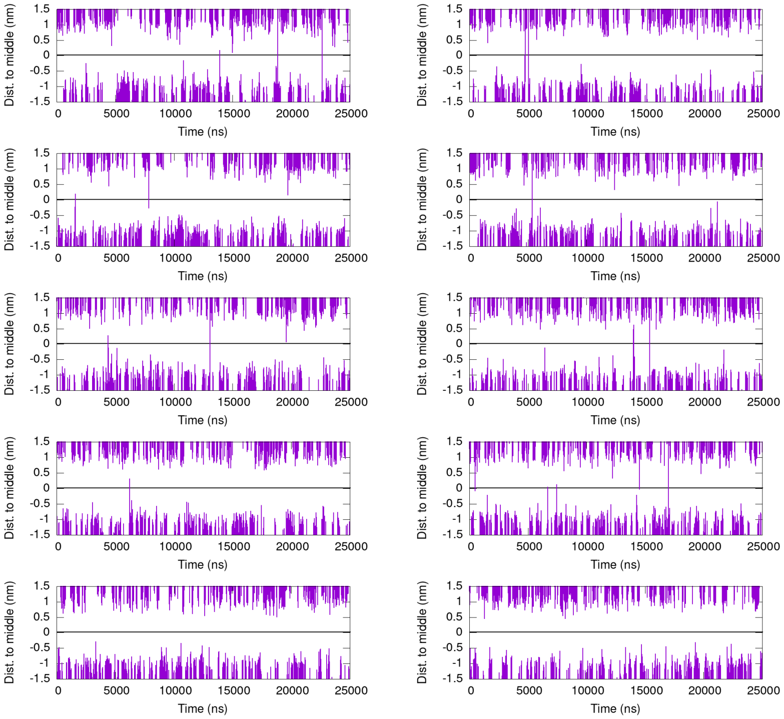

Figure S9: Distance of the COM of each of the 10 SALBT drug molecules to the middle of the membrane along the simulation for one of the replicas in the presence of β2AR. *Flip-flops* can be observed when the distance changes from positive to negative values or vice versa.

### 8 Flat-bottom potential for SALBT simulations: Number of flip-flops, density plot, and distance of the ligands to the middle of the membrane

Since the density distributions of SALBT were not showing the occurrence of *flip-flops* due to the diffusion of the drug molecules through the periodic boundary conditions (pbc), a flat-bottom potential was applied in the same simulations performed with and without the protein ($\Delta$R = 3.5 nm, 1000 kJ/mol, and $\Delta$R = 6.0 nm, 10 kJ/mol, respectively). The potential is centered in the membrane middle, at the same time that the positions of the tail beads of the membrane were also restrained ($\Delta$R = 0.2 nm, 20 kJ/mol in both simulation setups).

Table S2: Number of *flip-flops* of SALBT molecules calculated for five replicas (25 µs simulation time each) of two different system setups; two of them including β2AR and two of them without. For these two different setups, two different simulations were run; a completely unbiased one and one including a flat-bottomed potential to avoid molecules overcoming pbc.

| | *Only POPC* | | *β2AR and POPC* | | | |
|---|---|---|---|---|---|---|
| | | | *Unbiased* | | *Flat-bottom potential* | |
| | *Unbiased* | *Flat-bottom Potential* | *Total* | *On β2AR* | *Total* | *On β2AR* |
| *Replica 1* | 1 ↓ 2 ↑ | 6 ↓ 2 ↑ | 4 ↓ 4 ↑ | 1 ↓ 2 ↑ | 6 ↓ 1 ↑ | 4 ↓ 1 ↑ |
| *Replica 2* | 1 ↓ 1 ↑ | 6 ↓ 1 ↑ | 3 ↓ 2 ↑ | 3 ↓ 0 ↑ | 6 ↓ 1 ↑ | 2 ↓ 1 ↑ |
| *Replica 3* | 3 ↓ 2 ↑ | 4 ↓ 3 ↑ | 4 ↓ 3 ↑ | 1 ↓ 2 ↑ | 6 ↓ 3 ↑ | 5 ↓ 2 ↑ |
| *Replica 4* | 2 ↓ 1 ↑ | 5 ↓ 3 ↑ | 1 ↓ 0 ↑ | 0 ↓ 0 ↑ | 5 ↓ 2 ↑ | 2 ↓ 1 ↑ |
| *Replica 5* | 3 ↓ 2 ↑ | 4 ↓ 0 ↑ | 2 ↓ 1 ↑ | 2 ↓ 0 ↑ | 11 ↓ 5 ↑ | 7 ↓ 3 ↑ |
| **Total** | 10 ↓ 8 ↑ | 25 ↓ 9 ↑ | 14 ↓ 10 ↑ | 7 ↓ 4 ↑ | 34 ↓ 9 ↑ | 20 ↓ 8 ↑ |

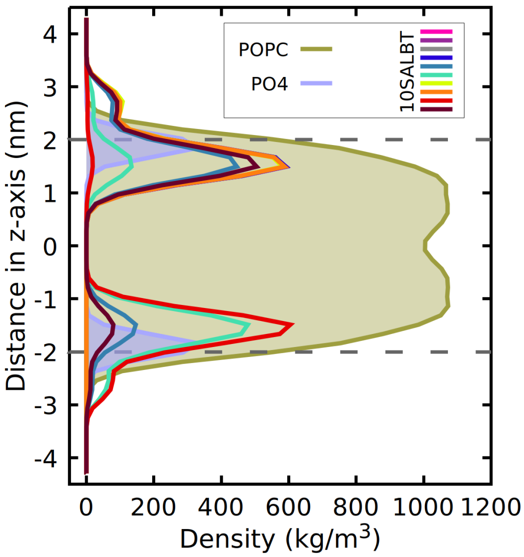

Figure S10: Density distribution of ten SALBT molecules, the POPC membrane, and the lipids' PO4 beads depicted along the membrane normal (z-axis) from a 25 µs simulation trajectory. The intracellular region corresponds to negative z-values; the extracellular region to positive z-values. The densities of the SALBT molecules are scaled by a factor of 100.

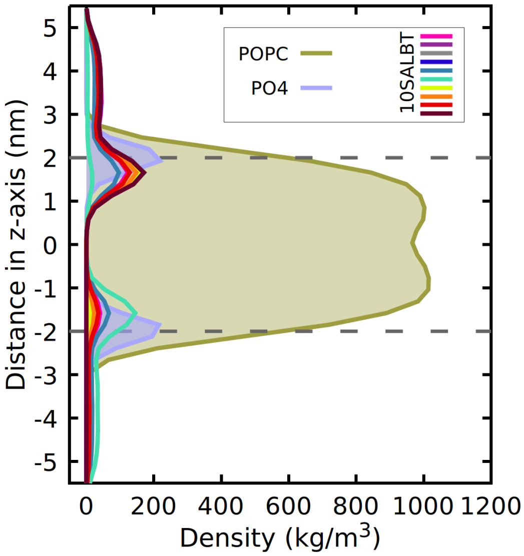

Figure S11: Density distribution of ten SALBT molecules, the POPC membrane, and the lipids' PO4 beads depicted along the membrane normal (z-axis) from a 25 µs simulation trajectory. The intracellular region corresponds to negative z-values; the extracellular region to positive z-values. The densities of the SALBT molecules are scaled by a factor of 100.

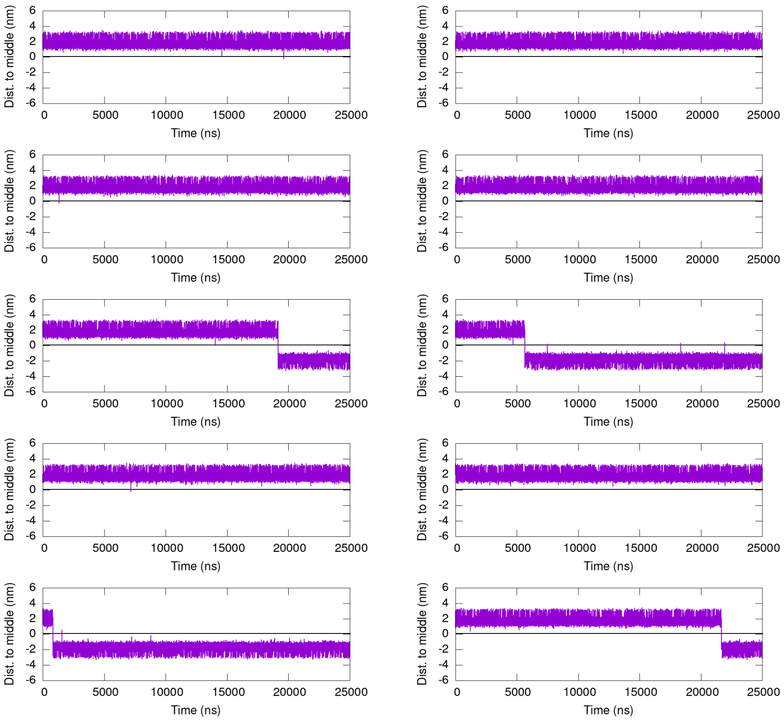

Figure S12: Distance of the COM of each of the 10 SALBT drug molecules to the middle of the membrane along the simulation for one of the replicas in the absence of β2AR. *Flip-flops* can be observed when the distance changes from positive to negative values or vice versa.

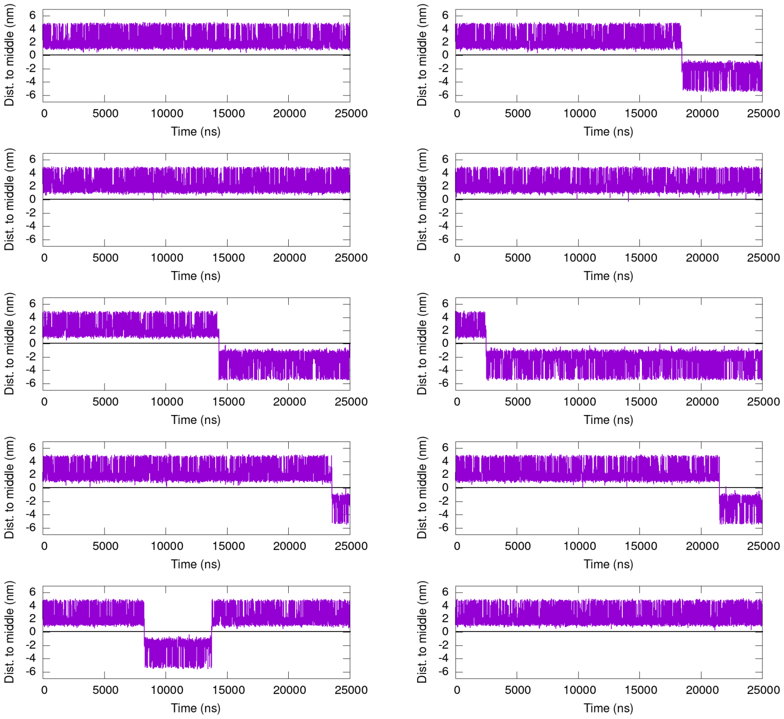

Figure S13: Distance of the COM of each of the 10 SALBT drug molecules to the middle of the membrane along the simulation for one of the replicas in the presence of β2AR. *Flip-flops* can be observed when the distance changes from positive to negative values or vice versa.

#### 9 Flip-flop localization on β2AR

##### Flip-flop analysis for SALMT

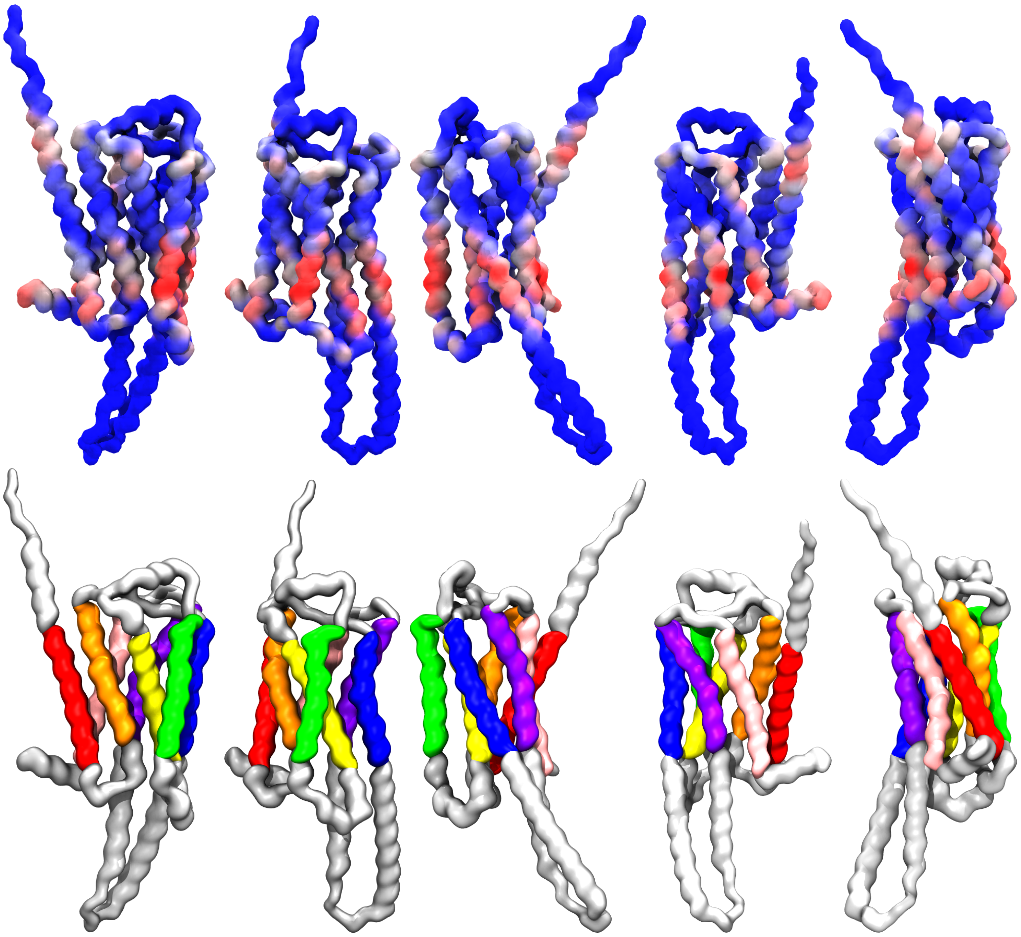

Figure S14: Five different viewpoints of β2AR are shown in two different color codes. The upper figures are colored according to the number of contacts of each residue with SALMT. Corresponding the bluest regions to the lowest amount of contacts and getting paler and then red as the number of contacts increases. The lower figures display the seven transmembrane domains previously mentioned employing the same color code as in Figure 3.

#### Flip-flop analysis for SALBT

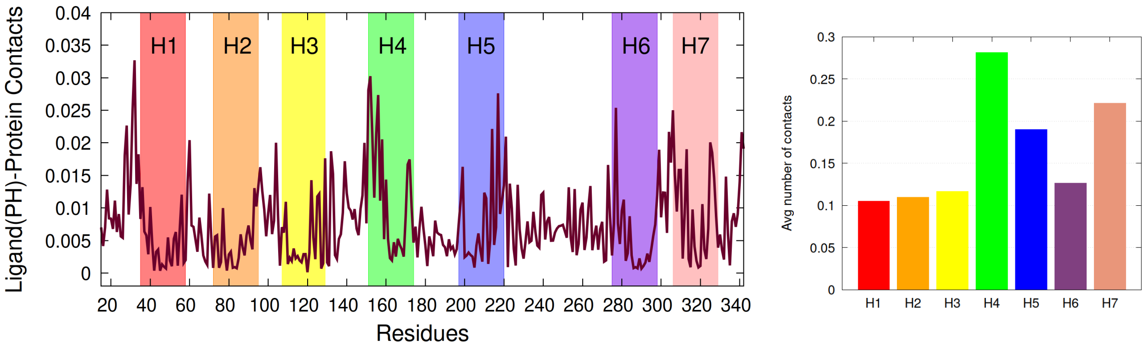

Figure S15: (Left) Average number of contacts between the headgroup of SALBT with the individual residues of β2AR for the five replicas (25 μs simulation time each). The colored rectangles indicate the seven transmembrane domains (H1-H7) of β2AR. (Right) Average number of contacts of the transmembrane domains (H1-H7) with the SALBT headgroup averaged over the five replicas.

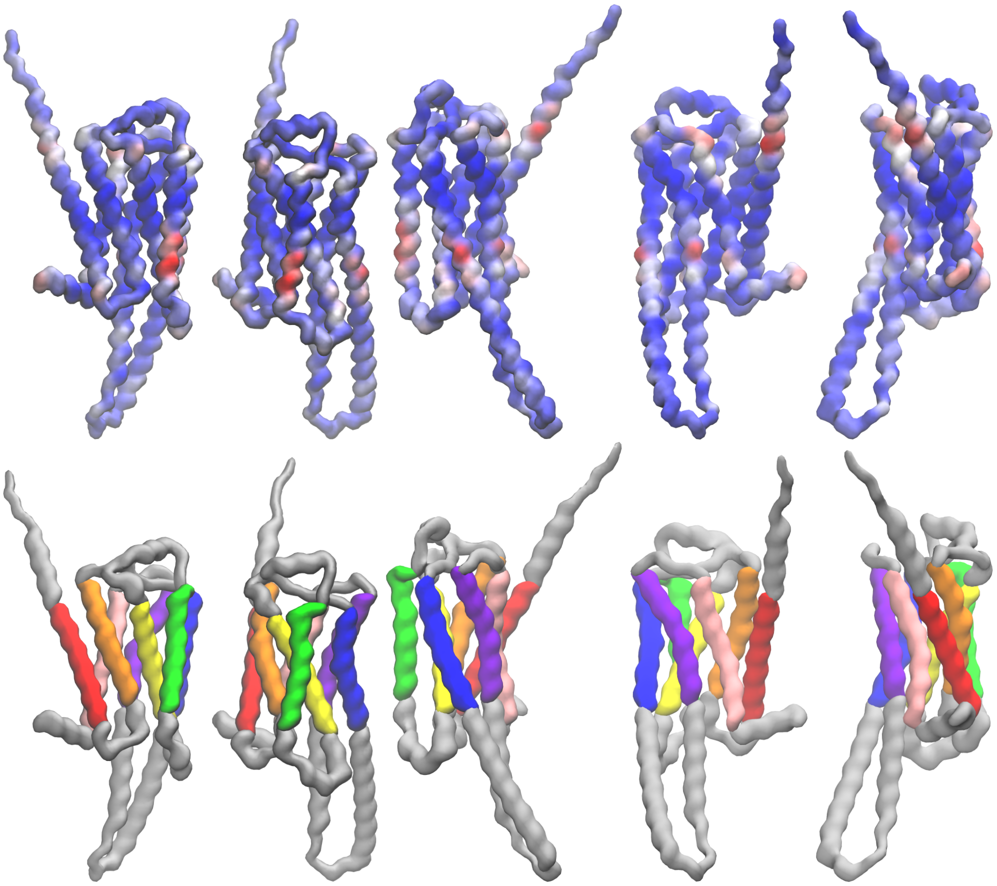

Figure S16: Five different viewpoints of β2AR are shown in two different color codes. The upper figures are colored according to the number of contacts of each residue with SALBT. Corresponding the bluest regions to the lowest amount of contacts and getting paler and then red as the number of contacts increases. The lower figures display the seven transmembrane domains previously mentioned employing the same color code as in Figure S15.

### 10 US convergence tests and histograms

The convergence tests show the PMF computed for increasing junks of 200 ns starting from the first 200 ns, when the system equilibration is considered to be finished.

The histograms show the bins of each window in different colours and the sum of them is displayed with a black line.

**US of SALMT from the water phase to the center of POPC membrane and from the center to the opposite edge of the bilayer (PO4 NC3)**

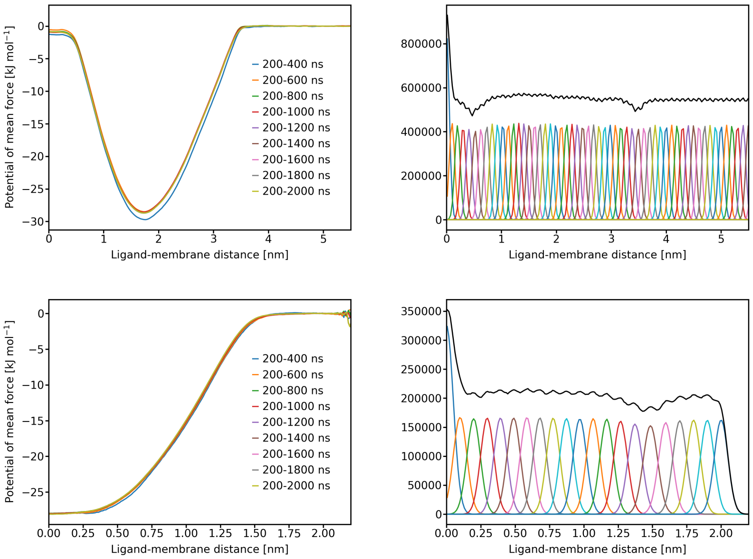

Figure S17: (Top, left) Convergence test and (top, right) histogram of the US of a SALMT molecule pulled from the water phase towards the center of the POPC membrane. (59 windows, every 0.1 nm, 2 µs per window). (Bottom, left) Convergence test and (bottom, right) histogram of the US when the SALMT molecule is pulled from the center to the opposite edge of the bilayer where the PO4 and NC3 beads of the lower leaflet are (21 windows, every 0.1 nm, 2 µs per window).

**US of SALBT from the water phase to the center of POPC membrane and from the center to the opposite edge of the bilayer (PO4 NC3)**

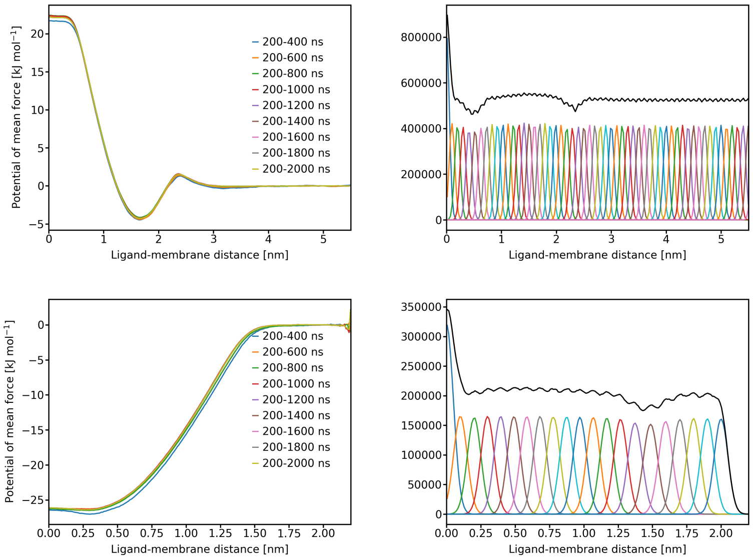

Figure S18: (Top, left) Convergence test and (top, right) histogram of the US of a SALBT molecule pulled from the water phase towards the center of the POPC membrane. (57 windows, every 0.1 nm, 2 µs per window). (Bottom, left) Convergence test and (bottom, right) histogram of the US when the SALBT molecule is pulled from the center to the opposite edge of the bilayer where the PO4 and NC3 beads of the lower leaflet are (21 windows, every 0.1 nm, 2 µs per window).

**US of SALMT along the β2AR protein embedded in a POPC membrane**

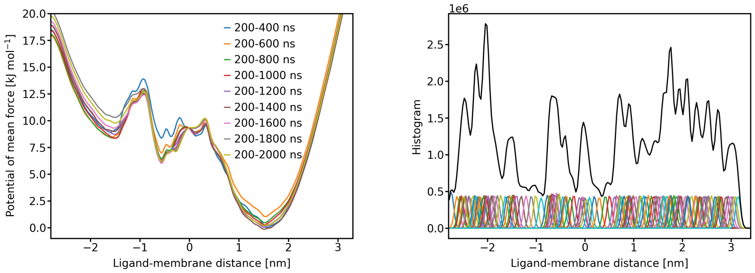

Figure S19: (Left) Convergence test and (right) histogram of the US performed by extracting the snapshots from the unbiased trajectory of a SALMT molecule *flip-flopping* along β2AR around the H4 domain (80 windows, 2 µs per window). And the extension of the sampling by pulling the molecule at both edges, inside and outside (18 + 32 windows, 2 µs per window).

#### US of SALBT along the β2AR protein embedded in a POPC membrane

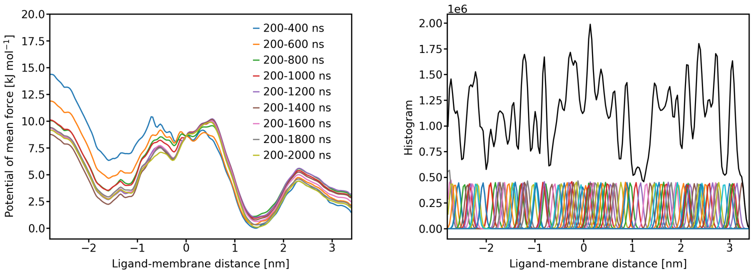

Figure S20: (Left) Convergence test and (right) histogram of the US performed by extracting the snapshots from the unbiased trajectory of a SALBT molecule *flip-flopping* along β2AR around the H4 domain (77 windows, 2 µs per window). And the extension of the sampling by pulling the molecule at both edges, inside and outside (15 + 29 windows, 2 µs per window).

#### US of SALBT in SALMT trajectory along the β2AR protein

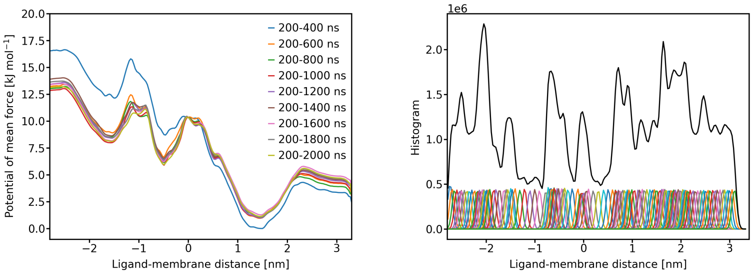

Figure S21: (Left) Convergence test and (right) histogram of the US performed by substituting SALMT for SALBT in the windows from the *flip-flop* trajectory. (123 windows, 2 µs per window).

#### US of SALMT in SALBT trajectory along the β2AR protein

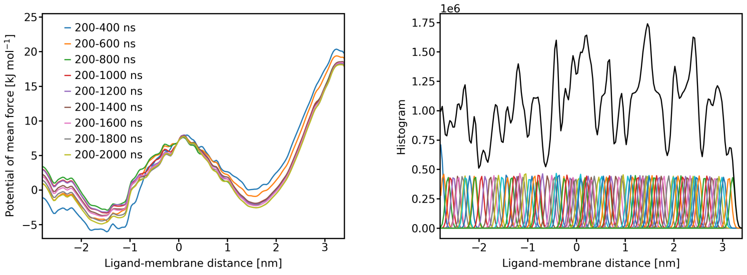

Figure S22: (Left) Convergence test and (right) histogram of the US performed by substituting SALBT for SALMT in the windows from the *flip-flop* trajectory. (113 windows, 2 µs per window).

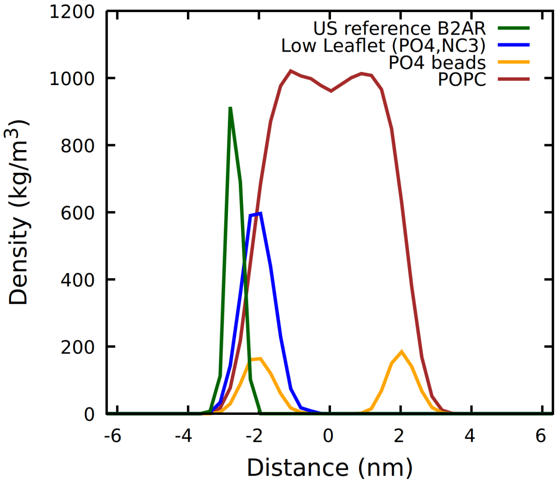

Figure S23: Density distribution of the beads taken as reference for the reaction coordinate of the US along the β2AR protein (US reference β2AR), the lipids' beads of the lower leaflet employed in the US of only the POPC membrane (Low Leaflet (PO4,NC3)), the whole POPC membrane density plot, and the lipids' PO4 beads depicted along the membrane normal (z-axis).

### 11 Dasatinib and baricitinib density plots

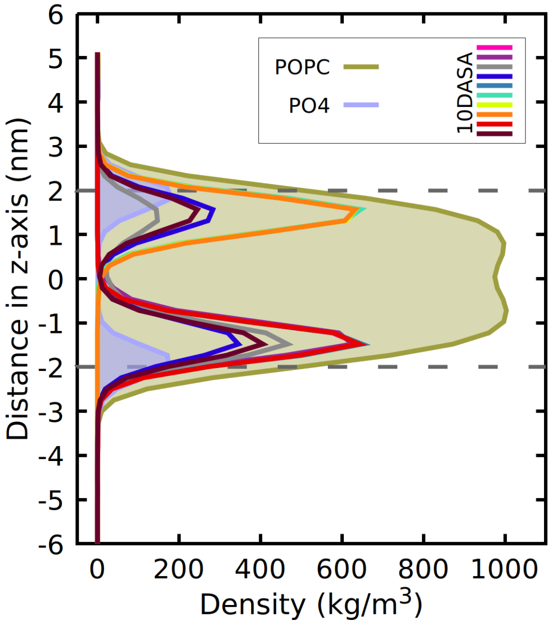

Figure S24: Density distribution of ten dasatinib (DASA) molecules, the POPC membrane, and the lipids' PO4 beads depicted along the membrane normal (z-axis) from a 25 µs simulation trajectory. The intracellular region corresponds to negative z-values; the extracellular region to positive z-values. The densities of the DASA molecules are scaled by a factor of 100.

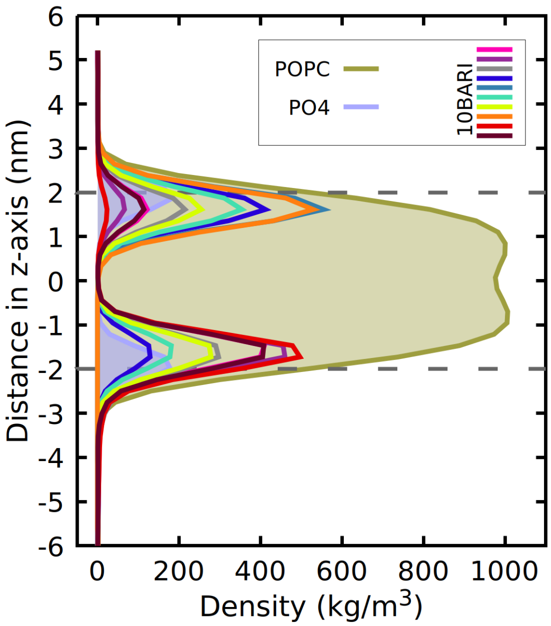

Figure S25: Density distribution of ten baricitinib (BARI) molecules, the POPC membrane, and the lipids' PO4 beads depicted along the membrane normal (z-axis) from a 25 µs simulation trajectory. The intracellular region corresponds to negative z-values; the extracellular region to positive z-values. The densities of the BARI molecules are scaled by a factor of 100.
